# Pan-Viral Conformational Landscapes of Frameshifting Elements Reveal Length-Dependent Plasticity and Antisense-Driven Structural Reprogramming

**DOI:** 10.64898/2026.05.13.724818

**Authors:** Kushal Adhikary, Abhishek Dey

## Abstract

Programmed ribosomal frameshifting (PRF) is an essential strategy used by many RNA viruses to expand their coding capacity within compact genomes. This process is governed by frameshifting elements (FSEs), specialized RNA structures that regulate translation through dynamic secondary and tertiary conformations. While the structural adaptability of the SARS-CoV-2 FSE has been extensively characterized, the conformational landscapes of FSEs across other pathogenic viruses remain poorly understood. Here, we present a comparative structural analysis of FSEs from Japanese Encephalitis Virus (JEV), West Nile Virus (WNV), Hepatitis C Virus (HCV), and Human Immunodeficiency Virus (HIV) using integrative computational modeling and molecular simulations. Our analysis reveals previously uncharacterized, virus-specific conformational ensembles, alongside a conserved core architecture that exhibits pronounced length-dependent structural plasticity. Extension of flanking sequences induces substantial conformational rearrangements, highlighting the role of sequence context in shaping FSE topology and potentially modulating PRF efficiency. Importantly, we demonstrate that antisense oligonucleotide (ASO) binding can reprogram FSE architectures, disrupting native structural motifs and stabilizing alternative conformations with altered thermodynamic stability. Collectively, these findings establish viral FSEs as dynamic RNA ensembles governed by sequence context and external interactions, and position ASO-mediated structural perturbation as a promising strategy for modulating frameshifting and viral gene expression.

## Introduction

Viruses utilise various strategies to manipulate host protein synthesis machinery and regulate the translation of their own proteins within the host cells. Programmed ribosomal frameshifting (PRF) is one of the biological phenomena employed by the RNA viruses to translate alternate proteins from their overlapping open reading frames [1,2]. This mechanism involves the altered movement of translating ribosomes on the viral genome, which effectively changes the translating frame as the ribosome now reads a different codon sequence, producing an alternative protein [3,4]. PRF is mainly of two types: +1 PRF, where the translating ribosomes read a new frame by skipping one nucleotide in the 3’ direction [3], and -1 PRF, where translating ribosomes first pause and then move back by one nucleotide towards the 5’ end to start translating from a new frame [5]. In addition to the viral proteome expansion, this mechanism also empowers the virus to fine-tune its gene expression, thus providing an additional layer to its regulatory mechanism [6]. A further, less common type of programmed ribosomal frameshifting (PRF) also involves +1 and -2 shifts in the reading frame by translating ribosomes [7].

Viral ribosomal frameshifting is regulated by different mechanisms involving different RNA regulatory elements present in the viral genome [7]. -1 PRF is guided by three such regulatory regions, which involve a slippery sequence and an RNA structural element, both joined to each other by the spacer region. The slippery sequence is the region that makes the translating ribosomes slip, as the codons of the slippery sequence are rarely associated with tRNA [5]. This enables an efficient -1PRF in the virus. The downstream region of this slippery sequence is connected to an RNA secondary structure element through a spacer. Presence of a particular RNA structure enhances the effect of the slippery site by stalling the ribosomes and making it backtrack by -1 nucleotide to change its reading frame [1] and translate multiple proteins [8]. -1 PRF is employed by Severe Acute Respiratory Syndrome Coronavirus-2 (SARS-CoV-2) to translate alternate proteins from its overlapping ORFs (ORF1a and ORF1b) [8–12].

Due to its immense therapeutic significance, the SARS-CoV-2 frameshifting element (FSE) has become one of the most extensively studied frameshifting regions, highlighting the multiple architectural folds this RNA can adopt. 6.9 Å cryo-EM structure of SARS-CoV-2 FSE identified it as a pseudoknotted structure appearing as a λ shaped conformation with the 5’ end being highly flexible [9]. However, a recent study involving RNA chemical modification followed by mutational profiling and deconvoluting algorithms has determined multiple interchangeable topologies of SARS-CoV-2 FSE [10,11]. These ensembles are in equilibrium and critical for efficient ribosomal frameshifting, as the structure-stabilizing mutants of SARS-CoV-2 FSE fail to frameshift effectively [11]. A parallel computational study also revealed the dynamic conformational landscape of the SARS-CoV-2 FSE in response to the inclusion of increasing numbers of nucleotides both upstream and downstream of the element [12].

Akin to SARS-CoV-2, many flaviviruses and retroviruses also display ribosomal frameshifting elements in their genome, crucial for viral proliferation and pathogenicity [13–19]. Flaviviruses are viruses with ∼10kb long positive-sense RNA encoding many structural and non-structural proteins. The major representatives of flaviviruses include Japanese Encephalitis Virus (JEV), Hepatitis C virus (HCV), and West Nile Virus (WNV), amongst others. All these viruses are known to translate alternative proteins from overlapping open reading frames through PRF. Japanese Encephalitis Virus (JEV) employs PRF to translate non-structural NS1’ protein essential for viral replication, assembly and host immune evasion [15]. Likewise, WNV also executes -1 PRF to translate the NS1 protein from the NS2A gene [16,20]. Though NS1 does not contribute to WNV replication and packaging, it contributes to WNV virulence by increasing neuroinvasiveness [15]. In Hepatitis C Virus, ribosome frameshift in -2/+1 frame causes the production of 17 kDa alternative frameshifting (F) protein from the overlapping reading frame present at the 5’ end of HCV ORF [17,21,22]. HCV F proteins induces apoptosis in human dendritic cells, thereby weakening host immune responses and facilitating viral persistence [23], and its antibodies are found to be circulating in the patient samples with acute HCV infection [24,25]. Retroviruses are RNA viruses which generate a copy of DNA from their RNA genome upon infection, followed by the integration of viral DNA into the host genome [26]. Human Immunodeficiency Virus (HIV) is one of the retroviruses known to display PRF to translate gag-pol polyprotein present at the 3’ end of *gag* ORF in the virus [27]. Gag protein is a viral structural protein essential for viral packaging [28], while Pol polyprotein gives rise to non-structural HIV proteins like integrase, reverse transcriptase, and protease [29]. The expression level of Gag protein in the Gag-Pol polyprotein is a tightly regulated process and is mediated by -1 PRF [30]. This ratio is crucial for maintaining viral structure and infectivity, and any deviations will inhibit HIV proliferation [31].

Although numerous studies have investigated the consequences of programmed ribosomal frameshifting (PRF) in viruses, structural analyses of their frameshift elements (FSEs) remain largely unexplored. Previous RNA SHAPE studies on various West Nile Virus (WNV) strains have revealed conformational plasticity in their FSEs, attributed to sequence variability at the 3′ end. However, comprehensive insights into the global conformational landscape of FSEs from these viruses, particularly in the context of their additional genomic nucleotide sequences, are still lacking. Using a combinatorial computational approach and tools (simRNA [32], trRosettaRNA [33], and AlphaFold3 [34]), molecular simulations, and principal component analysis (PCA), we have deconvoluted the conformational ensembles of RNA frameshift elements (FSEs) from JEV, WNV, HCV, and HIV, both globally and in the presence of various antisense oligonucleotides (ASOs). Despite JEV, WNV, and HCV belonging to the same viral family, their FSEs adopt distinct tertiary conformations. Notably, FSE topologies remained consistent across different computational algorithms, underscoring the robustness of our determined models. We observed that adding nucleotides upstream, downstream, or on both sides of the core FSE sequence induces conformational shifts, highlighting the dynamic nature of FSEs. Furthermore, we characterised ASO-induced structural remodelling of FSEs, underscoring the feasibility of modulating frameshifting elements through targeted antisense interactions.

## Results

### Sequence variability in the frameshifting regions

To investigate the conformation of FSEs in JEV, WNV, HCV, and HIV, we initially selected 85-nucleotide-long FSE regions for analysis (Fig. 1A). This sequence length was chosen based on previous studies focused on viral frameshifting regions [9,10,11,12, 35]. To assess conservation among the FSE regions of different viruses, we performed multiple sequence alignments using Clustal Omega. The alignment revealed notable sequence variability across the 85-nt FSEs of the various viruses (Fig. 1B). Among these, the JEV and WNV FSEs showed the highest sequence similarity (∼85%). In contrast, the HCV FSE shared lower similarity with both JEV (47%) and WNV (41%) (Fig. 1B). Interestingly, the HIV FSE exhibited greater sequence identity with JEV (58%) and WNV (56%) than with HCV (46%) (Fig. 1B).

**Figure 1.**
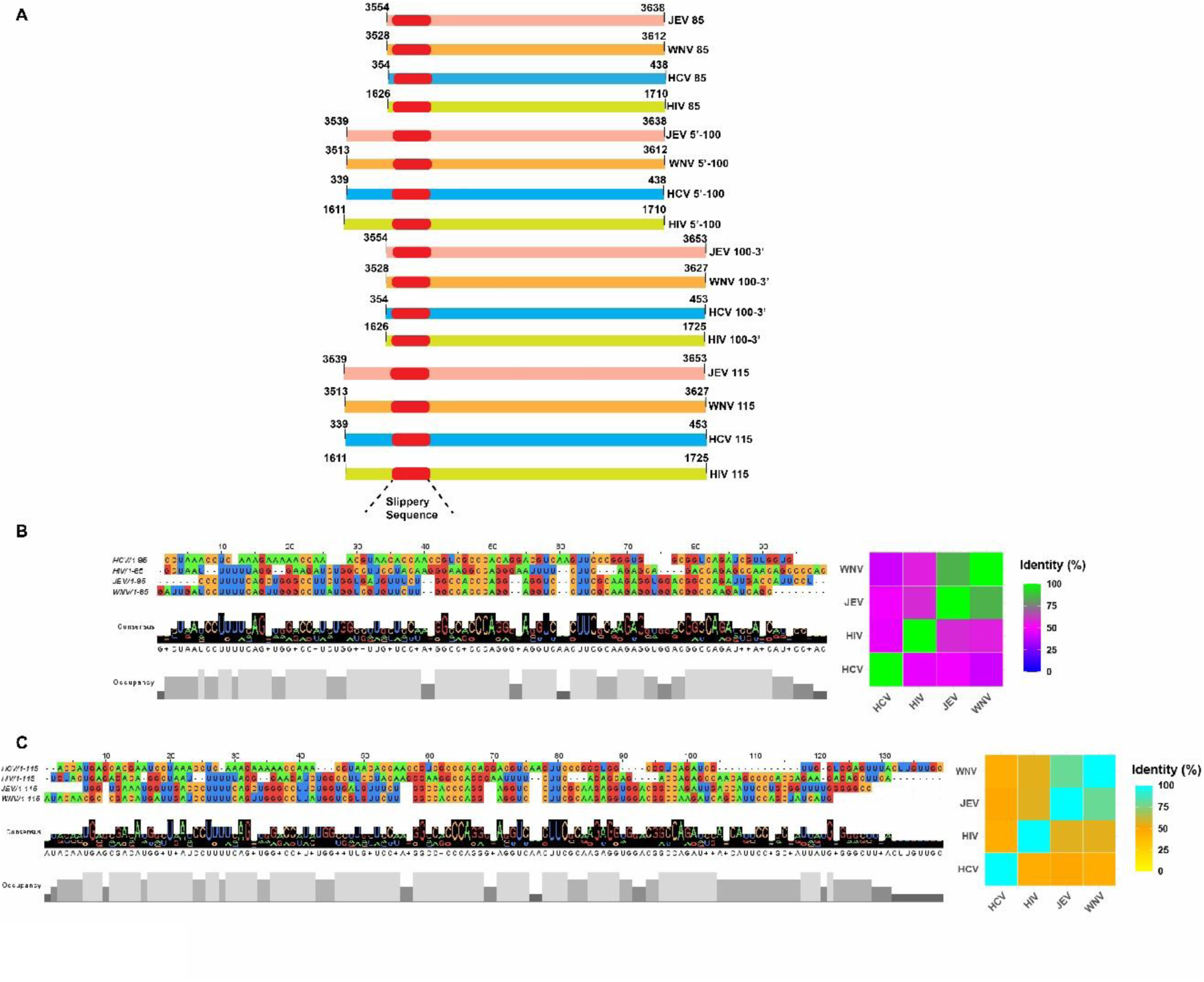
**(A)** Schematic representation of frameshifting region (including slippery site in red color) in the genomic RNA of Japanese Encephalitis Virus (JEV), West Nile Virus (WNV), Hepatitis C Virus (HCV), and Human Immunodeficiency Virus (HIV) each for 85nt, 100-3’ nt, 5’-100nt, and 115 nt. (**B)** Sequence alignment of 85nt FSE region along with its per cent identity matrix in the form of a heat map **(C)** Sequence alignment of 115nt FSE region along with its per cent identity matrix in the form of a heat map. Sequence alignment of 85nt and 115nt FSE constructs shows higher similarity between the FSEs of JEV, WNV and HIV.

Despite these differences, several nucleotide positions were highly conserved across the viruses. Notably, the slippery sequence was highly conserved in the FSEs of JEV, WNV, and HIV, with a preference for poly-U tracts (Fig. 1B). In contrast, HCV displayed a poly-A slippery sequence. Conserved residues across all FSEs included A at positions 50, 55, and 86; U at positions 10, 58, 63, and 64; G at positions 16, 43, 70, 72, 80, and 85; and C at positions 48, 49, 65, and 82 (Fig. 1B).

To evaluate the influence of flanking sequences on FSE architecture, we generated extended constructs by adding 15 nucleotides upstream (5’-100), downstream (100-3’), or at both ends (115-nt total) of the 85-nt FSEs (Fig. 1A). Consistent with the 85-nt sequence alignment results, the 115-nt FSEs of JEV and WNV remained highly similar (∼80%), followed by moderate similarity with the HIV 115-nt FSE (55%) (Fig. 1C). HCV’s 115-nt FSE continued to show relatively low similarity with those of JEV, WNV, and HIV (47%, 40%, and 46%, respectively) (Fig. 1C). Similar trends were observed for the 5’-100 and 100-3’ FSE constructs, mirroring the conservation patterns seen in the 85-nt and 115-nt alignments (Fig.S1). Notably, a greater degree of divergence was observed between the FSE regions of HCV and WNV (Fig. 1 and Fig. S1).

### 85nt FSEs adopt a distinct yet similar architecture

Next, we applied various folding algorithms to predict the conformation of 85-nt FSE regions from all four viruses. Initially, we used IPknot to determine their secondary structures. Among the four FSEs, those from JEV and HCV adopted pseudoknotted conformations, while the FSEs of WNV and HIV formed unknotted stem-loop helical structures (Table S1). To further explore their structural organisation, we modelled the three-dimensional (3D) architecture of each FSE using different methods, including simRNA, trRosettaRNA, and AlphaFold3 and verified the predicted models using molecular simulation studies. Despite employing distinct folding algorithms, each software consistently predicted a similar conformation for the FSEs, with subtle structural differences unique to each virus. (Fig. 2).

**Figure 2.**
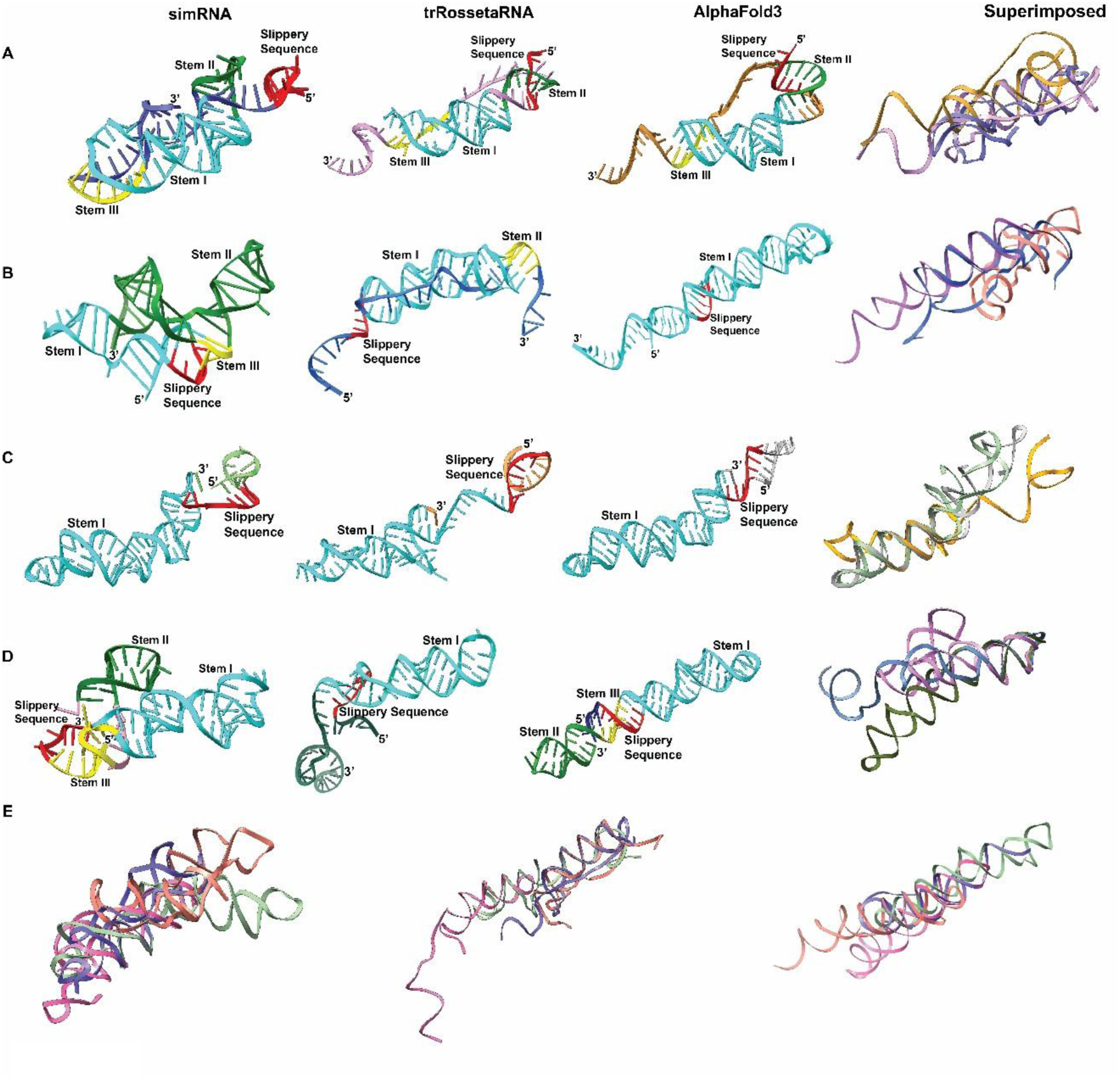
Tertiary structure prediction of 85nt FSE using simRNA, trRossetaRNA, and AlphaFold3 for **(A)** JEV, **(B)** WNV, **(C)** HCV, and **(D)** HIV. The last column represents the superimposition of each viral FSE model generated through simRNA, trRosettaRNA, and AlphaFold3. **(E)** Superimposition of different viral FSEs architecture generated through simRNA, trRosettaRNA, and AlphFold3.

For JEV, simRNA predicts that the frameshift element (FSE) adopts a pseudoknotted structure comprising three distinct stems—Stem I (Cyan), Stem II (forest green) and Stem III (yellow), with Stem II and Stem III forming two separate pseudoknots (Fig. 2A). In contrast, trRosettaRNA predicted an unknotted stem-loop helical conformation (Fig. 2A). However, a similar arrangement observed in the AlphaFold3 model (Fig. 2A), shows Stem III pairing with the apical loop of Stem I to form a pseudoknot, and the slippery site engaged in base-pairing with Stem II, creating a simple stem-loop motif. The overall model obtained from different algorithms is further reinforced by several non-canonical interactions (Table S2). Despite these differences, all three models generated for the 85nt JEV FSE exhibit a root-mean-square deviation (RMSD) of approximately 1 Å, indicating overall structural similarity with subtle conformational variations (Fig. 2A).

For the 85-nt WNV FSE, the predicted knotted structures generated by simRNA, and trRosettaRNA shows distinct pseudoknot configurations (Fig. 2B). The pseudoknot predicted by simRNA features base-pairing interactions between the slippery sequence and Stem III, forming a knotted structure (Fig. 2B). In contrast, trRosettaRNA predicted pseudoknot formations involving interactions between the apical loop of Stem I and Stem III, while the slippery sequence remained single-stranded in these models (Fig. 2B). Surprisingly, contrary to simRNA and trRosettaRNA, AlphaFold3 did not identified any pseudoknotted architecture (Fig. 2B). Rather, it determined the WNV 85nt FSE as a single stem-loop helical conformation (Fig. 2B). Despite these structural differences, a comparative analysis indicates substantial conformational overlap among the models, with RMSD values ranging from 0.8 Å to 1.2 Å (Fig. 2B).

Although IPknot predicted a pseudoknotted structure for the 85-nt HCV FSE, simRNA, trRosettaRNA, and AlphaFold3 consistently folded the above construct into a two-way stem-loop junction (Fig. 2C). Among these, the models generated by simRNA and AlphaFold3 displayed highly similar conformations, with an RMSD of approximately 0.75 Å (Fig. 2C). Notably, in both of these models, the slippery sequence pairs with the 5′ end of the FSE to form a small stem-loop helix (Fig. 2C). In contrast, the model predicted by trRosettaRNA exhibited distinct base-pairing patterns compared to simRNA and AlphaFold3. Specifically, nucleotides U72–C78 form an internal bulge in the trRosettaRNA model, resulting in a higher RMSD of around 1.0 Å when compared with the other two models (Fig. 2C).

Interestingly, the 85-nt HIV FSE adopted three distinct conformations when modelled using simRNA, trRosettaRNA, and AlphaFold3 (Fig. 2D). simRNA predicted a two-stem-loop structure with the slippery sequence appeared to interacting with the 5′ end, forming an extra stem-loop helix characterized by triple base-pairing involving A61–U3–A12 (Fig. 2D). In contrast, trRosettaRNA predicted a single stem-loop (Stem I), with the 3′ end remaining largely unstructured (Fig. 2D). Meanwhile, AlphaFold3 modelled the FSE as a two-way stem-loop junction (a dumbbell shape structure with one more extended stem-loop arm at 5’ end and one shorter stem-loop arm at 3’ end), where the slippery sequence interacted with Stem III nucleotides similar to simRNA predictions at the base of Stem I, and the 3′ end base-paired to form Stem II (Fig. 2D). Due to these significant structural differences, the RMSD values among the three models were found to be ≥1.0 Å (Fig. 2D).

To further evaluate the architectural similarities among FSEs from different viruses, we superimposed their 3D models, generated using respective RNA folding algorithms and visualised the pairwise RMSD values as a heatmap (Fig. 2E). Based on the simRNA predictions, the 85-nt FSE of JEV showed a high degree of structural similarity (RMSD ≤ 1.0 Å) to the FSEs of WNV, HCV, and HIV (Fig. 2E). However, the HCV FSE exhibited a distinct conformation compared to the WNV FSE, with an RMSD > 1.0 Å. Similar structural trends were observed with trRosettaRNA-derived models, except that the HIV FSE displayed a different conformation relative to the JEV and WNV FSEs (Fig. 2E). In contrast, AlphaFold3 generated a markedly divergent model for the HCV FSE, with substantial differences (RMSD ≥ 1.0 Å) from the FSEs of JEV, WNV, and even HCV itself when compared across algorithms.

To further analyze the FSE folding, tertiary RNA models were converted into secondary structure representations using x3DNA-DSSR [49]. Comparative arc plots were subsequently generated using the R-Chie web server [49] to assess conformational concordance among structures predicted by simRNA, trRosettaRNA, and AlphaFold3. Arc-plot analysis revealed that the frameshifting elements (FSEs) of JEV, HCV, and HIV adopt comparable conformation in simRNA- and AlphaFold3-derived models (Fig. 3). Notably, simRNA predicted two pseudoknots within the JEV FSE, whereas AlphaFold3 resolved a single pseudoknot (Fig. 3A). For most viral FSEs, simRNA and AlphaFold3 models exhibited similar architectures, with WNV representing a notable exception (Fig. 3B). Both simRNA and trRosettaRNA predicted knotted conformations for the WNV FSE (Fig. 3B); however, simRNA specifically identified a kissing-loop–mediated knotted architecture (Fig. 3B). In contrast, AlphaFold3 predicted an unknotted conformation for the WNV FSE, distinct from both simRNA- and trRosettaRNA-derived models (Fig. 3B). Collectively, comparative arc-plot analysis complements the tertiary-structure-based assessment and indicates substantial concordance between simRNA- and AlphaFold3-generated models for all the 4 viral FSEs (Fig. 3).

**Figure 3.**
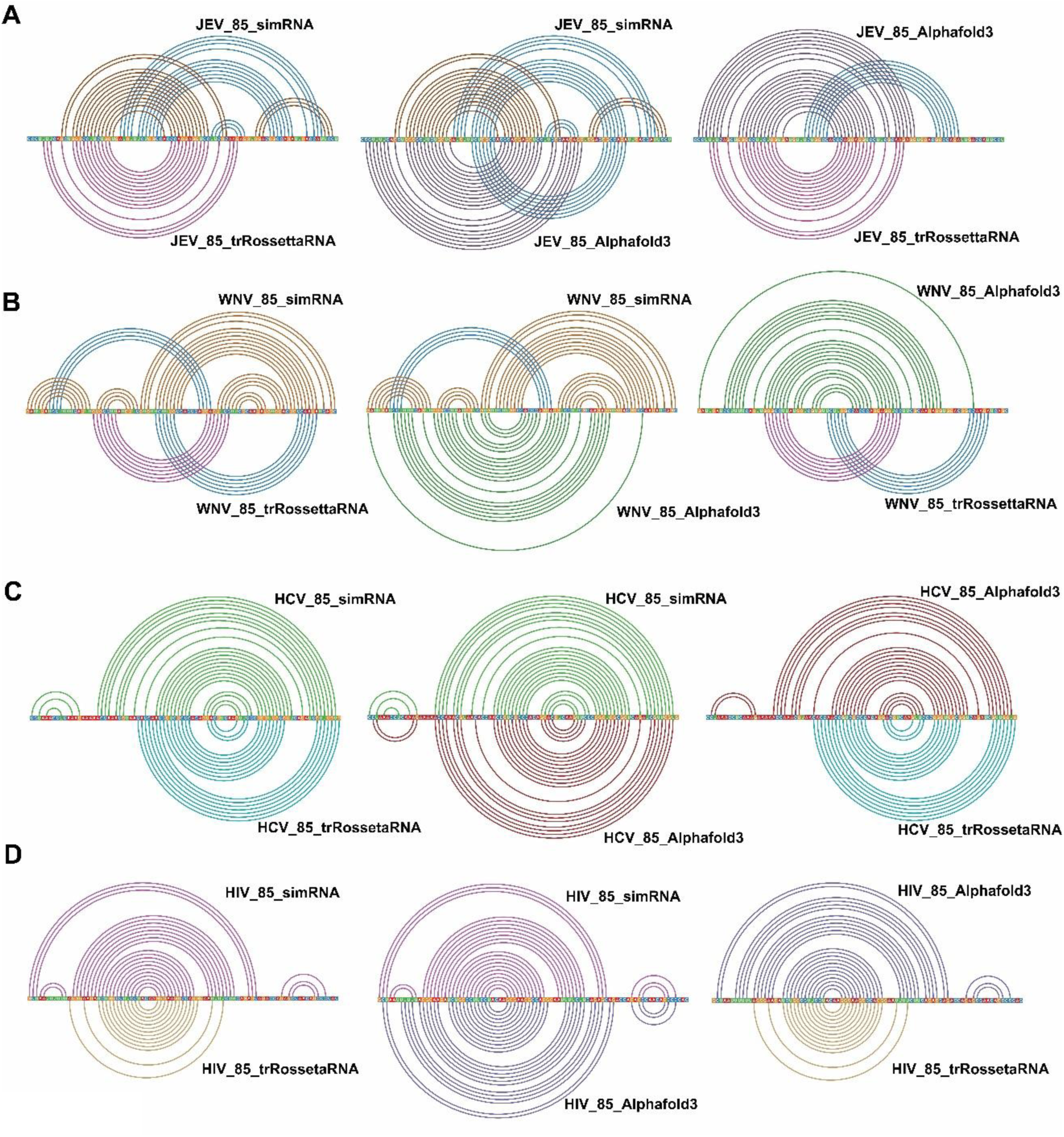
Comparative arc plots between the 85 nt FSE models generated using simRNA, trRosettaRNA, and AlphaFold3 for **(A)** JEV, **(B)** WNV, **(C)** HCV, **(D)** HIV. Comparative analysis largely similar folding between the models generated from simRNA and AlphaFold3 when compared to trRosettaRNA generated models.

We further validated the simRNA- and AlphaFold3-derived models using molecular dynamics simulations. Across systems, the simulations largely preserved the global folds of the viral FSEs, yielding conformations comparable to the respective starting models and exhibiting low RMSD values (Fig. 4). In most cases, simRNA-derived structures showed closer agreement with their simulated counterparts. Notably, both JEV and WNV FSEs consistently adopted pseudoknotted conformations in the simRNA predictions and their corresponding simulations (Fig. 4A, 4B). Similarly, the HCV FSE models generated by simRNA and their corresponding simulated structures were found to be in close concordance (Fig. 4C). In contrast, simulations initiated from the simRNA-derived HIV FSE model converged toward a more open conformation, characterized by reduced base pairing and diminished stabilization of helical elements relative to the initial simRNA model (Fig. 4D). Comparable trends were also observed for simulations initiated from AlphaFold3-predicted HIV FSE structures (Fig. 4D).

**Figure 4.**
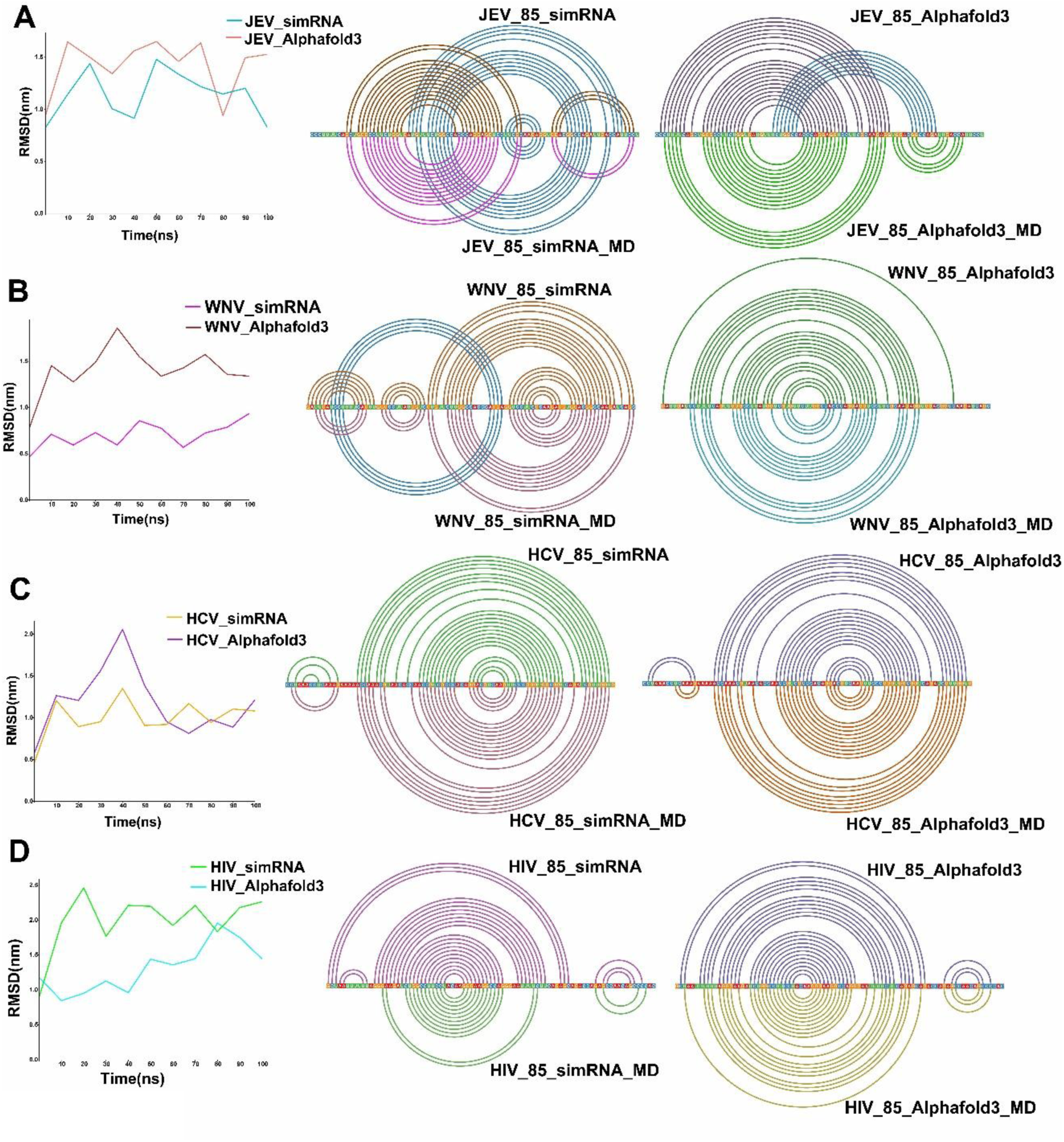
Root Mean Square Deviation (RMSD) plots and comparative arc plot analysis between the models generated from simRNA and AlphaFold3 and their simulated models generated from GROMACS 2025.4 for **(A)** JEV FSE 85nt, **(B)** WNV FSE 85nt, **(C)** HCV FSE 85nt, and **(D)** HIV FSE 85nt. The analysis represents identical base-pairing interactions between the models generated from simRNA and AlphaFold3 with their simulated counterparts. Colored arcs represent base-pairing interactions, enabling visualization of secondary-structure, including pseudoknots and long-range contacts.

Simulations of AlphaFold3-predicted FSEs from WNV, HCV, and HIV largely retained conformations similar to their respective predicted models (Fig. 4B, 4C, 4D). An exception was observed for the JEV FSE AlphaFold3 predicted model, where simulations unexpectedly favored an unknotted architecture (Fig. 4A). Despite this disparity, the core JEV FSE maintained an identical stem–loop helical architecture with identical base pairing in both the AlphaFold3-predicted structure and its simulated ensemble (Fig. 4A).

To understand the conformational landscape of each viral FSEs we performed time-resolved principal component analysis of simRNA and AlphaFold 3 generated simulated models (Fig. 5). This analysis revealed distinct conformational sampling patterns across viruses and modeling strategies (Fig. 5). In all systems, trajectories progressively expand from early (0–1 ns) (red dots) to intermediate (1-99 ns) (orange dots) to late (99–100 ns) (blue dots) phase, indicating continuous exploration of the conformational space.

**Figure 5.**
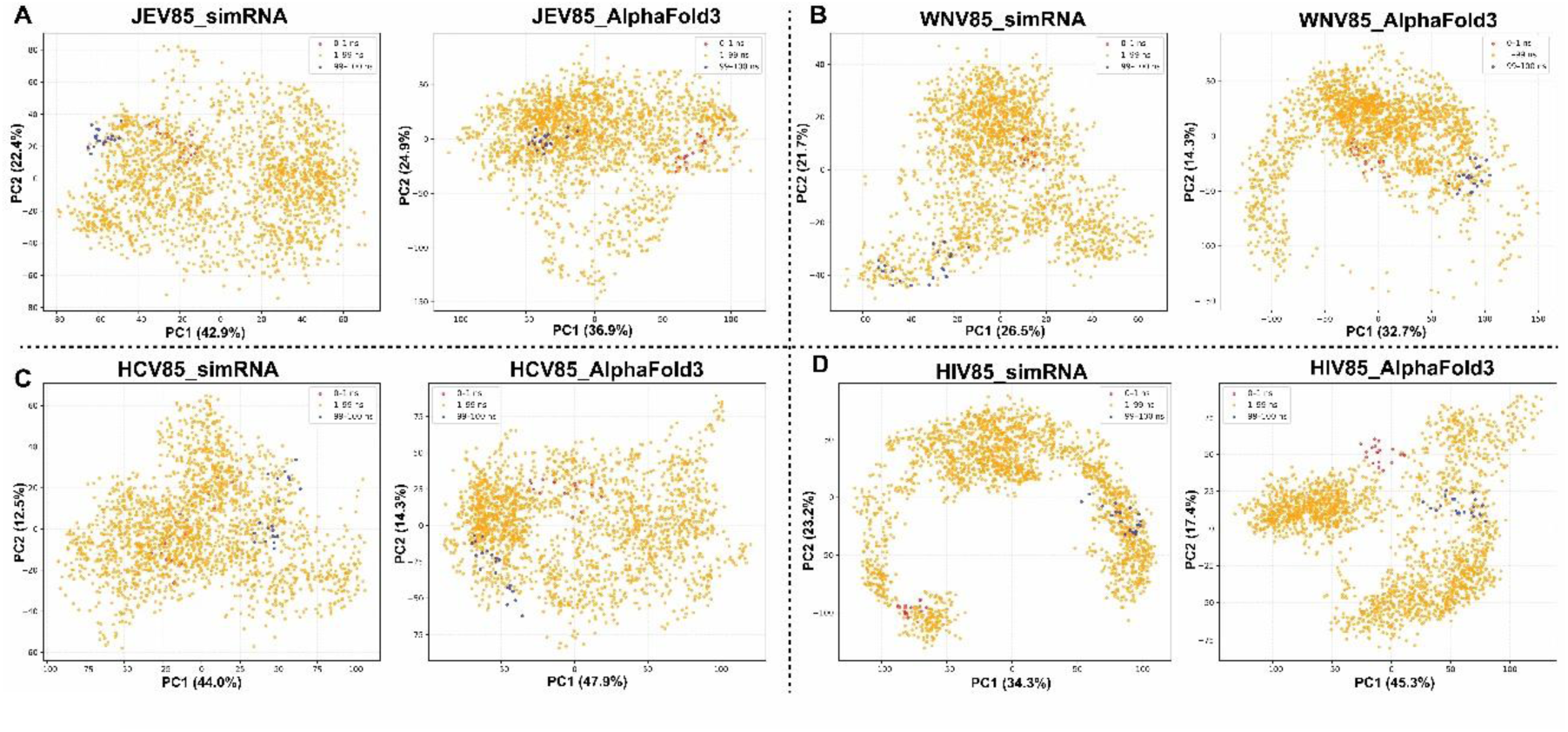
Time-resolved PCA reveals conformational ensembles of 85-nt viral FSEs modeled using simRNA and AlphaFold3 for **(A)** JEV, **(B)** WNV, **(C)** HCV, and **(D)** HIV. simRNA structures display broader dispersion and multiple basins, indicating higher heterogeneity, whereas AlphaFold3 models show tighter clustering, suggesting restricted sampling and stabilization in defined conformational states.

For JEV (Fig. 5A), simRNA-derived structures exhibit a broad and dispersed distribution across space, suggesting the presence of multiple conformational topography. In contrast, AlphaFold3 models show relatively more compact clustering with partial spreading at later time points, indicating moderate conformational flexibility but reduced heterogeneity compared to simRNA. In WNV (Fig. 5B), AlphaFold3 models display an extended, arc-like distribution, reflecting structured yet directional conformational transitions. simRNA ensembles, however, are more diffusely distributed with less defined trajectories, indicating stochastic sampling of multiple structural states.

For HCV (Fig. 5C), both simRNA and AlphaFold3 models show substantial dispersion, but AlphaFold3-derived ensembles dispersed across occupy a wider x-axis, suggesting enhanced large-scale structural rearrangements. simRNA models, while heterogeneous, appear comparatively centralized, indicating constrained exploration relative to AlphaFold3. In HIV (Fig. 5D), simRNA trajectories form a pronounced ring-like distribution, indicative of highly dynamic conformational transitions between transient states. AlphaFold3 models, in contrast, show partitioned clustering with distinct conformational states, suggesting stabilization in free energy minima rather than continuous transitions.

Overall, while modelling of the 85-nt FSEs from JEV, WNV, HCV, and HIV using these three algorithms generally yielded similar topologies, subtle yet notable differences were observed. Since, simRNA and AlphaFold3 yielded comparable FSE topologies across all viruses, we selected simRNA and AlphaFold3 generated models for subsequent analyses.

### Flanking nucleotides induces notable structural modifications in JEV and WNV, HCV, and HIV FSEs

To evaluate how flanking nucleotides influence the overall conformation of the viral frameshift elements (FSEs), we extended the JEV FSE sequence by adding 15 nucleotides either to the 5′ end, the 3′ end, or to both ends, and subsequently folded these longer constructs using simRNA and AlphaFold3 (Fig. 1A). Folding the 5′-extended FSE with simRNA resulted in a more compact, circular architecture (Fig. 6A, Fig. S5A). In contrast, adding 15 nucleotides at the 3′ end yielded a more elongated FSE conformation (Fig. 6A, Fig. S3A). Notably, the addition of 15 nucleotides to both ends (resulting in a 115-nt construct) led to a significant transformation, with the JEV FSE adopting a distinct three-way junction (3WJ) fold (Fig. 6A, Fig. S3A). AlphaFold3 predictions revealed that the 5′-100, 100-3′, and full 115-nt JEV constructs formed compact, circular architecture (Fig. S3B). Additionally, AlphaFold3 also predicted pseudoknots stabilised through kissing loop interactions in all the longer JEV constructs (Fig. S2A, Fig. S3B). In the 115-nt construct, the 5′ end (in stem-loop helical form) base-pairs with the 3′ end to generate a second pseudoknot (Fig. S2A). Consistent with simRNA results, AlphaFold3-predicted structures for JEV FSE also revealed notable differences in their overall architecture (Fig. S2A, Fig. S3A, and S3B).

**Figure 6.**
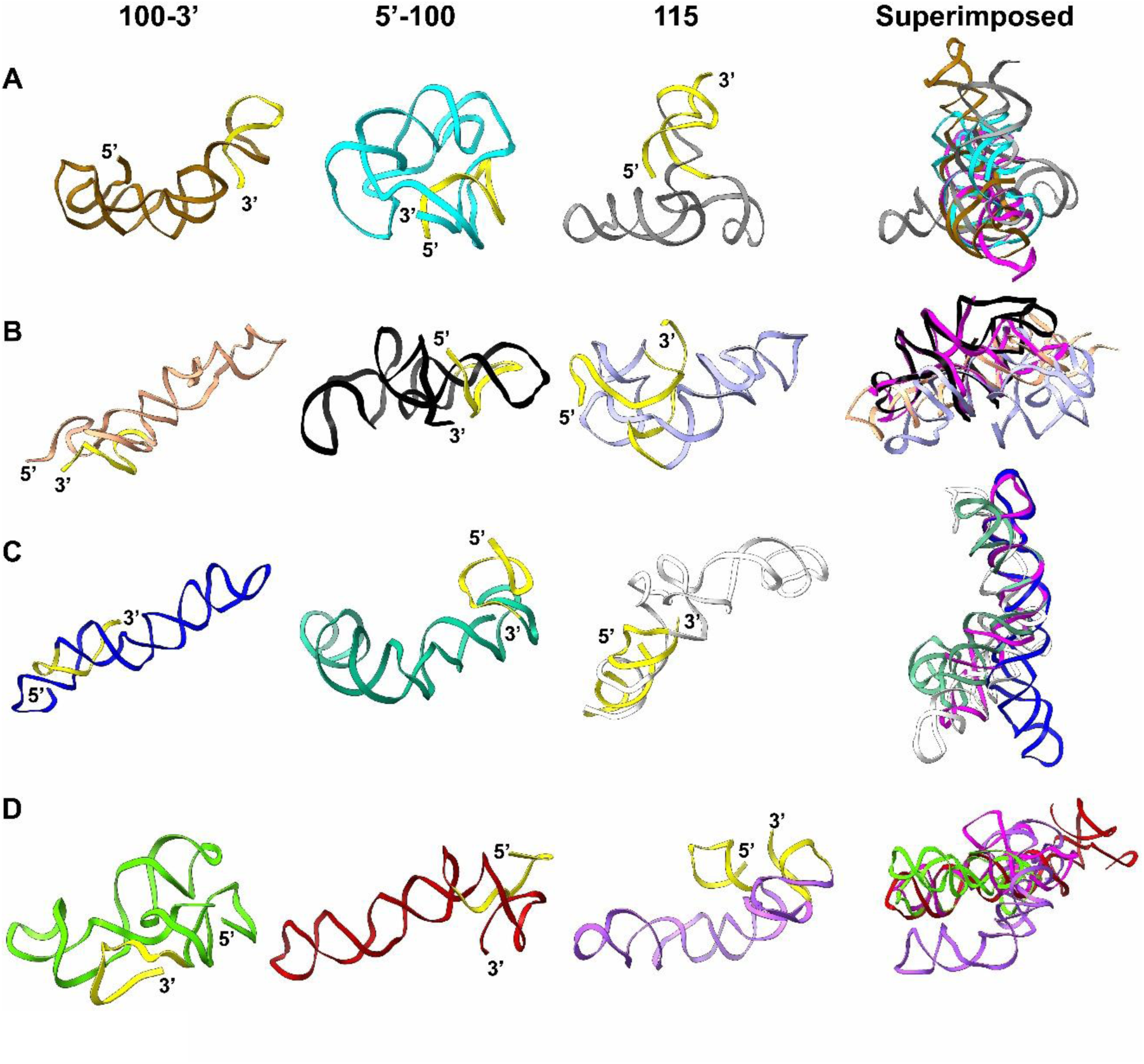
SimRNA-generated tertiary structure model of larger FSE constructs (100-3’, 5’-100, and 115nt) for **(A)** JEV, **(B)** WNV, **(C)** HCV, and **(D)** HIV. Additional nucleotide in each construct are shown in yellow color. The last column represents the comparative folding analysis between longer FSE constructs and the 85nt FSE of respective viruses (pink colour). Overall, length-dependent conformational flexibility was observed in the longer FSE constructs of all the viruses compared to the 85nt FSE.

Comparative arc-plot analysis of simRNA- and AlphaFold3-derived JEV FSE models reveals a highly conserved core base-pairing across the minimal 85-nt FSE and all extended constructs, indicating robust preservation of the frameshifting core architecture (Fig. S4). Extension of the FSE sequence introduces additional long-range base-pairing interactions, particularly in the 5′-100 and 115-nt constructs, leading to the formation of extra pseudoknots and enhanced stabilization through 5′–3′ interactions (Fig. S4). In contrast, the 3′-100-nt construct forms knotted conformations but largely lacks long-range contacts (Fig. S4). Notably, both prediction platforms show consistent trends, underscoring that sequence extension modulates global architecture without perturbing the conserved FSE core.

A similar analysis was performed on the frameshift elements (FSEs) of WNV, HCV, and HIV using constructs of varying lengths. The WNV FSE 5’-100nt construct, modelled using simRNA, adopts a structure comparable to that of the 85-nt WNV FSE (Fig. 6B, Fig. S3C). However, the additional nucleotides at the 5’ end form a stem-loop structure, whose loop participates in a pseudoknot stabilised by three triple base-pairing interactions (Fig. 6B, Table S2). The 100–3’ construct adopts an elongated stem-loop helix conformation, different from knotted 85nt WNV FSE structure (Fig. 6B, Fig. S3C). Meanwhile, the 115-nt WNV FSE folds into a compact pseudoknotted structure (Fig. 6B, Fig. S3C). Notably, this pseudoknot differs from the one observed in the 85-nt structure (Fig. 6B). AlphaFold3 predicts that the 5′-100nt construct of the WNVFSE adopts an elongated stem-loop helix conformation devoid of any pseudoknot (Fig. S2B, Fig. S3D). This conformation is identical to that of the AlphaFold3 predicted 85nt WNV FSE structure (Fig. 2B). Similarly, the 100–3′nt WNV FSE construct folds into an elongated two-way junction stem-loop helix (Fig. S2B, Fig. S3D). Intriguingly, when additional nucleotides were incorporated at both ends to generate a 115nt construct, AlphaFold3 predicted a compact, circular structure featuring a pseudoknot (Fig. S2B, Fig. S3D). This configuration is markedly distinct from the extended, unknotted stem-loop conformation observed in the 85nt construct (Fig. S3C, S3D).

Comparative arc-plot analysis shows that WNV FSE constructs adopt multiple long-range interaction patterns, with clear conformation-dependent differences between simRNA- and AlphaFold3-derived models (Fig. S5). While both approaches capture the core interaction framework of the FSE, simRNA predictions display greater conformational diversity and redistribution of long-range contacts across the 5′-100, 3′-100, and 115 nt constructs (Fig. S5). In contrast, AlphaFold3 models appear more compact, particularly in longer constructs (Fig. S5). Overall, these comparisons highlight pronounced conformational plasticity of the WNV FSE and suggest that the FSE sequence strongly influences the WNV FSE folding landscape.

Addition of 15 nucleotides at the 5′ end of the HCV frameshift element (FSE) (5′-100nt construct) results in a curved stem-loop junction without a pseudoknot, as predicted by simRNA (Fig. 6C, Fig. S3E). Similarly, appending 15 nucleotides at the 3′ end (100–3′nt construct) yields an elongated structure with a pseudoknot localised at one end (Fig. 6C, Fig. S3E). When 15 nucleotides are added to the 5′ and 3′ ends (115nt construct), simRNA predicts a two-way stem-loop helical conformation with a curved overall architecture (Fig. 6C, Fig. S3E). Overall, the extended HCV FSE conformations predicted by simRNA closely resemble the structure of the 85 nt HCV FSE (Fig. 6C, Fig. S3E). Consistent with simRNA predictions, the AlphaFold3 modelled the 5′-100nt HCV FSE construct as a curved stem-loop helical conformation (Fig. S2C, Fig. S3F). The 100–3′nt construct adopted an elongated stem-loop helical junction, forming a two-way junction with stem-loop helices positioned at opposite ends (Fig. S2C, Fig. S3F). A similar two-way stem-loop junction conformation was also observed in the AlphaFold3 prediction of the 115nt HCV FSE construct (Fig. S2C, Fig. S3F). In this model, one of the stem regions is formed through base pairing between nucleotides at the 5′ and 3′ termini (Fig. S2C, Fig. S3F). Overall, adding nucleotides at either end did not significantly alter HCV FSE architecture, which remained broadly consistent with the structure predicted for the 85nt construct by AlphaFold3 (Fig. S3E, S3F).

Arc-plot comparisons of HCV FSE constructs reveal a conserved core folding pattern across all lengths, with both simRNA and AlphaFold3 capturing the dominant long-range interactions characteristic of the HCV frameshifting element (Fig. S6). However, simRNA models exhibit greater heterogeneity, with multiple alternative interaction sets and pronounced rearrangements upon extending the sequence from 85 nt to 115 nt or altering nucleotide addition at either 5′ or 3′ regions. In contrast, AlphaFold3 predictions converge on more compact and uniform conformations, maintaining similar interaction architectures in core region with additional long-range interactions observed for 3’ and 115 nt constructs (Fig. S6). These observations indicate that, while the HCV FSE is structurally robust, its peripheral regions remain conformationally flexible and sensitive to sequence context strategy (Fig. S6).

While simRNA predicted a knotted structure for the 85-nt HIV FSE (Fig. 2D), extending the sequence by 15 nucleotides at the 5′ end resulted in an elongated single stem-loop helix structure (Fig. 6D, Fig. S3G). Similarly, adding 15 nucleotides at the 3′ end also produced a distinct, compact conformation featuring pseudoknots stabilised by two kissing-loop interactions, as predicted by simRNA (Fig. 6D, Fig. S3G). The longest construct comprising the core FSE region extended by 15 nucleotides at the 5′ and 3′ ends adopted a unique curved stem-loop helical architecture (Fig. 6D, Fig. S3G). While initial analysis suggested an absence of pseudoknots, closer inspection of the 3′ end of the 115-nt construct revealed a triple-helical motif indicative of a smaller pseudoknot at this end (Fig. 6D, Fig. S3G).

For the 5′-100nt construct of the HIV FSE, AlphaFold3 predicted an extended three-stem-loop helical structure arranged as a three-way junction (3WJ) (Fig. S2D, Fig. S3H). This conformation closely resembles the 85nt HIV FSE construct, also folded using AlphaFold3 (Fig. S2D, Fig. S3H). A similar 3WJ architecture was observed for the 100–3′nt construct (Fig. S2D, Fig. S3H). Interestingly, this conformation is distinct because it adopts a curved architecture, with the three stem-loop helices organised to form an overall λ-shaped structure (Fig. S3H). This λ-shaped fold contrasts with the extended conformation predicted for the original 85nt FSE construct by AlphaFold3 (Fig. S3H). Notably, when 15nt were added to both the 5′ and 3′ ends of the FSE, AlphaFold3 predicted a different structure featuring a central extended stem-loop flanked by two shorter stem-loops at either end, resulting in a bow-like architecture (Fig. S2D). Collectively, these findings highlight AlphaFold3’s ability to predict distinct structural conformations for various HIV FSE constructs, depending on sequence length and flanking regions (Fig. S3H).

Comparative arc plots of HIV FSE constructs reveal that the core frameshifting architecture is largely conserved across different sequence lengths and prediction methods (Fig. S7). simRNA models exhibit higher conformational variability with context-dependent long-range interactions upon 5′ or 3′ extensions, whereas AlphaFold3 predictions converge on more compact and consistent interaction patterns (Fig. S7). Notably, the extended constructs stabilize additional peripheral base-pairing without disrupting the central FSE core, highlighting intrinsic structural robustness coupled with sequence dependent conformational plasticity.

Similar to the 85-nt FSE constructs, the 115-nt FSE models generated using simRNA and AlphaFold3 were further validated through molecular dynamics simulations (Fig. 7). Simulation analysis indicates that the JEV FSE adopts a comparable global structure in both simRNA- and AlphaFold3-derived models, with AlphaFold3 simulations consistently preserving pseudoknot features (Fig. 7A). In the case of WNV, AlphaFold3 predicts pronounced knotted architectures, including kissing-loop interactions, whereas simRNA favors a distinct, predominantly unknotted topology (Fig. 7B). By contrast, HCV and HIV FSEs display broadly similar structural patterns across simRNA and AlphaFold3 models, with MD simulations largely maintaining the predicted secondary-structure features (Fig. 7C, 7D). Collectively, these results highlight both conserved and model-specific topological features among viral FSEs and further emphasize the complementary strengths of simRNA- and AlphaFold3-based structure prediction approaches.

**Figure 7.**
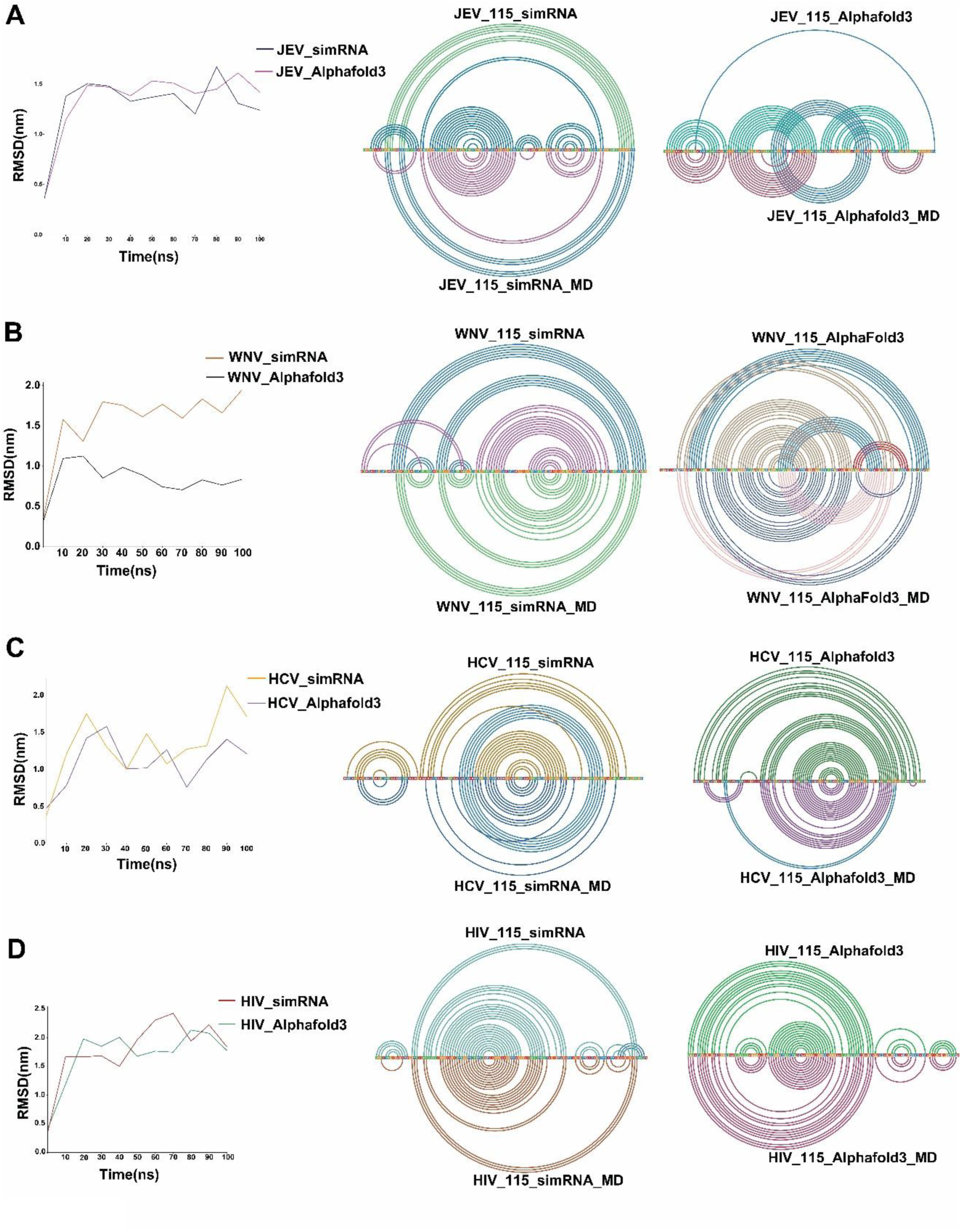
Comparative structural dynamics and arc-plot analysis of viral frameshifting elements (FSEs). Panels **A–D** show analyses for JEV, WNV, HCV, and HIV FSEs (115 nt), respectively. For each virus, the left subpanel depicts backbone RMSD profiles over 100 ns molecular dynamics (MD) simulations for simRNA- and AlphaFold3-derived models, indicating overall structural stability and convergence. The middle and right subpanels present arc plots derived for simRNA and AlphaFold3 models before and after MD refinement, respectively. Colored arcs represent base-pairing interactions, enabling visualization of secondary-structure, including pseudoknots and long-range contacts.

The PCA projections of the 115-nt viral FSE constructs reveal distinct conformational landscapes and time-dependent structural evolution across viruses and modelling approaches. For JEV (Fig. S8A), both simRNA and AlphaFold3-derived ensembles exhibit broad dispersion along both components, indicating substantial conformational heterogeneity. However, AlphaFold3 structures show more defined clustering, suggesting relatively stabilized conformational states as compared to simRNA. In WNV (Fig. S8B), the conformational space is comparatively more compact in AlphaFold3, with a dominant cluster centred near the origin, whereas simRNA displays a wider spread with distinct peripheral populations suggesting transient excursions into higher-energy conformations.

For HCV (Fig. S8C), both methods capture a highly heterogeneous ensemble, but simRNA exhibits a more diffuse and continuous distribution, indicating extensive conformational sampling. In contrast, AlphaFold3 shows partial segregation into subclusters, implying the presence of semi-stable structural intermediates. In HIV (Fig. S8D), a striking difference is observed where simRNA produces a curved, trajectory-like distribution, indicative of a directional conformational transition pathway over time. Conversely, AlphaFold3 yields a more scattered yet clustered landscape, lacking a well-defined conformational transition pathway path but highlighting multiple metastable states.

Overall, both approaches capture the flexible nature of viral FSEs. However, simRNA explores a broader and more continuous range of conformations, while AlphaFold3 identifies more distinct and stable structural states, indicating method-dependent differences in representing FSE dynamics.

### Antisense Oligonucleotide (ASO)-induces conformational changes in frameshift elements (FSEs)

To further assess the impact of ASOs on the conformational landscape of viral FSEs, we performed co-folding analyses using both 85-nt and 115-nt variants of each viral FSE. ASOs approximately 21 nucleotides in length, selected based on their low minimum free energy of duplex formation as predicted by OligoWalk (Table S3), were prioritised for this analysis. Co-folding was performed by providing the sequences of each ASO and the corresponding viral FSE into the RNAcofold web server (see methods section). The resulting dot-bracket notation (.dbn) files were used as hard constraints in simRNA to generate simulated tertiary structures of the ASO-FSE complexes. In parallel, we employed AlphaFold3 to independently predict the co-folded tertiary structures of the ASO and viral FSEs.

The ASO directed against the 85nt JEV construct disrupted the native pseudoknots by binding to its 5′ end, leading to the formation of a novel stem-loop structure at the 3′ end (Fig. 8A). This structural rearrangement resulted in an extended conformation of the JEV FSE (Fig. 8A). A similar extended conformation was observed when AlphaFold3 was used to co-model the ASO with the JEV FSE (Fig. 8B) where disruption of pseudoknot was observed, promoting the formation of a stem-loop helix at the 3′ end (Fig. 8B).

**Figure 8.**
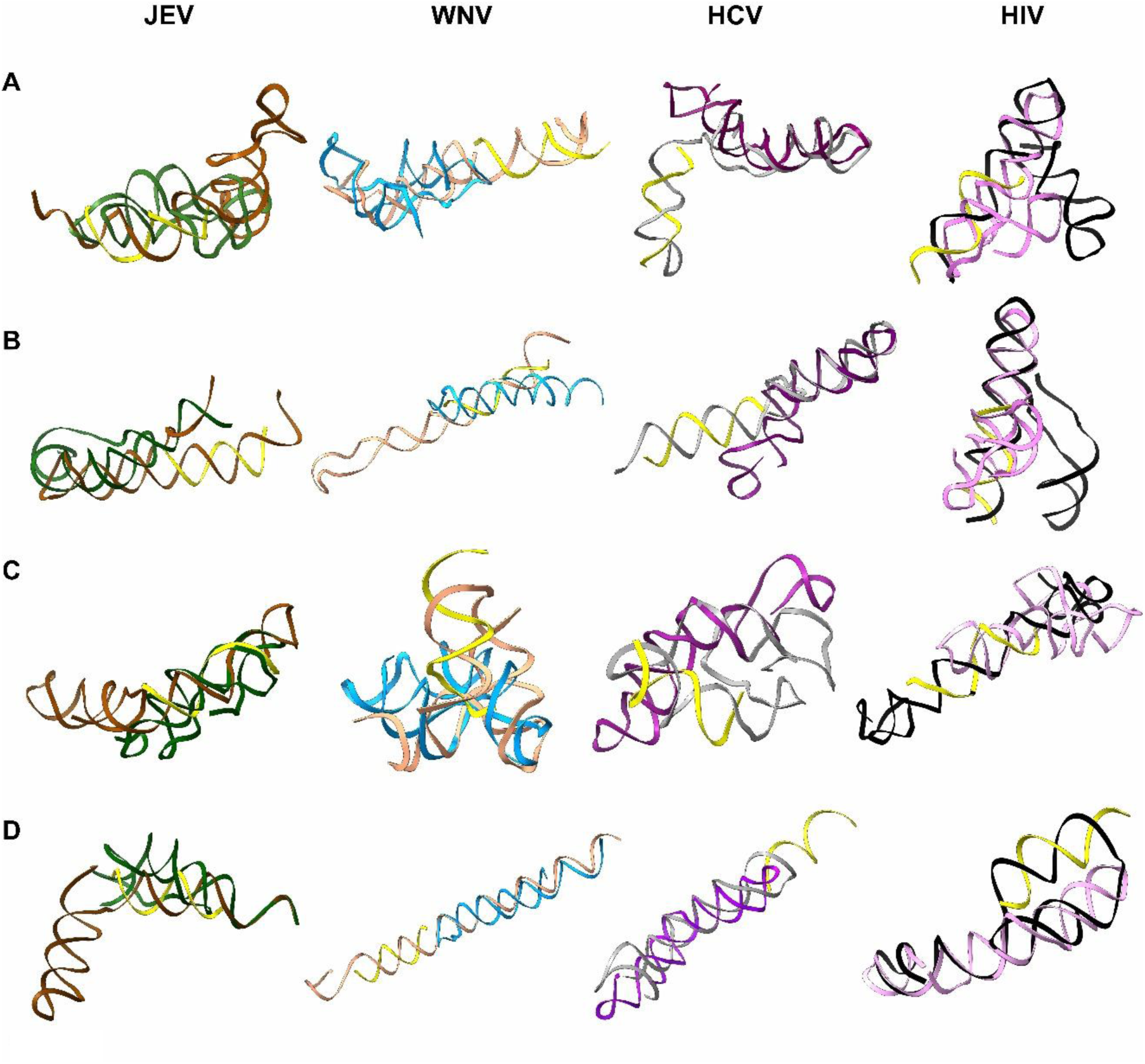
Co-folding analysis between ASO (yellow) generated through OligoWalk and binding to the 5’ end of 85nt FSEs of JEV, WNV, HCV, and HIV using **(A)** simRNA and **(B)** AlphaFold3. Co-folding analysis between ASO (yellow) generated through OligoWalk and binding to the central or 3’ end of 85nt FSEs of JEV, WNV, HCV, and HIV using **(C)** simRNA, **(D)** AlphaFold3. Superimposition over native 85nt FSE shows a significant alteration in the conformation of each viral FSE in the presence of ASO.

The ASO targeting the 85-nt WNV FSE disrupted its predicted knotted structure, as shown by simRNA modelling (Fig. 8A). This disruption led to the formation of a new stem-loop helix at the 3′ end of the WNV FSE (Fig. 8A). Although AlphaFold3 predicted an unknotted stem-loop helical conformation for the unbound 85-nt WNV FSE (Fig. 2B), co-folding analysis of the ASO with the WNV FSE using AlphaFold3 revealed a disruption in the base-pairing interactions within this helix (Fig. 8B). Specifically, the stem-loop was shortened in the co-folded ASO–WNV FSE complex compared to the unbound structure (Fig.7B).

For the 85-nucleotide HCV FSE-ASO complex, both simRNA and AlphaFold3, reveals disruption in the native stem-loop helix along with its shortening at the 3′ end in the co-modelled conformation (Fig. 8A, 8B). Binding of ASO to the 5′ end of the 85-nt HIV FSE led to the disruption of its native pseudoknotted conformation. simRNA predicted a λ-shaped structure, with one arm forming a duplex with the ASO at the 5′ end (Fig. 8A). A similar λ-shaped conformation of the HIV FSE in the presence of ASO was also predicted by AlphaFold3 (Fig. 8B), whereas the native HIV FSE was initially predicted to adopt a stem-loop helix structure (Fig. 2D).

OligoWalk generated a diverse set of ASOs targeting various regions of the viral FSE (Table S3). We next assessed the impact of ASOs on the overall architecture of the viral FSEs by co-folding to their respective targets but at different site. ASO binding at the 3’ end of the 85-nt JEV FSE disrupted its native structure, leading to the formation of an arc-like conformation, with the 3’ end participating in pseudoknot formation as predicted by simRNA (Fig. 8C). A similar arc-like architecture was also predicted by AlphaFold3; however, in this case, the 3’ end forms a simple stem-loop helix rather than engaging in a pseudoknot (Fig. 8D).

Binding of a second ASO to the 3′ end of the 85-nt WNV FSE led to a significant structural rearrangement, with simRNA predicting a shift from its native pseudoknotted conformation to a more compact, triangle-like architecture (Fig. 8C). In contrast, AlphaFold3 predicted a complete unwinding of the stem-loop helix within the WNV85 FSE upon ASO binding at the same site (Fig. 8D). Collectively, these findings suggest that ASO targeting the 3′ region of the WNV85 FSE causes a complete disruption of its native architecture.

OligoWalk also identified a second ASO that base-pairs with the central region of the HCV85 FSE (Table S3). Co-folding analysis using simRNA revealed a compact, circular conformation in which the 5′ and 3′ ends of the HCV85 FSE are brought into proximity (Fig. 8C) involving new base-pairing interactions. This structure represents a significant departure from the previously predicted extended conformation of the native HCV85 FSE (Fig. 2C). In contrast, AlphaFold3 predicted only subtle changes upon co-folding with the second ASO. While the overall conformation remained similar to the native extended architecture, minor variations were observed at both apical ends of the HCV85 FSE construct, with the 5’ end folding into an apical loop (Fig. 8D).

For the HIV85 FSE, OligoWalk also identified a second set of ASO targeting the central region of the element (Table S3). Co-folding analysis using simRNA in the presence of this ASO revealed substantial topological rearrangements within the HIV85 FSE. While simRNA initially predicted a pseudoknotted structure for the native HIV85 FSE, ASO binding led to an unfolding of this architecture, resulting in a more elongated conformation featuring two small stem-loop helices at either end (Fig. 8C). Co-folding analysis of the ASO with HIV85 FSE using AlphaFold3 revealed a dramatic structural transformation into a spatula-like conformation. In this conformation, the 5′ end of the RNA folds into a stem-loop helix, which subsequently folds back to form a pseudoknotted structure through base pairing with the RNA backbone. This back-folding generates a large central single-stranded RNA loop, serving as the binding site for the ASO, ultimately giving rise to the spatula-like architecture. This conformation markedly deviates from the stem-loop helix structure observed in the native HIV85 FSE (Fig. 8D, Fig. 2D). Similar analyses using larger FSE constructs (115 nt) from all the viruses revealed that binding of two independent ASOs at distinct sites within the viral FSE led to disruption of the native FSE architecture (Fig. S9).

To assess the thermodynamic stability and Boltzmann-weighted ensembles formed upon interaction of ASOs with their respective FSEs, we computed the base-pair partition function for each ASO–FSE complex. The partition function analysis consistently indicates stable heterodimer formation between ASOs and their target FSEs, as evidenced by a well-defined and dense interaction blocks in Region II, representing a highly stable and reproducible helical domain (Fig. S10, S11). In contrast, the more diffuse and fragmented interaction patterns observed in Region I, corresponding to intramolecular interactions within the viral FSEs, suggest the presence of multiple transient and alternative conformations (Fig. S10, S11). The stability of ASO–FSE interactions is further corroborated by the favorable (more negative) free energy values associated with these complexes (Table S4).

As a control, we generated scrambled versions of the ASOs and co-folded them with their respective FSEs. This analysis showed that, in the presence of scrambled ASOs, the FSEs retained their native structures without any significant structural disruption (Fig. S12). These results further validate the reliability of the predicted tertiary models of FSEs in complex with ASOs (Fig. S12). Overall, the findings from this study suggest that the architecture of viral FSEs can be effectively modulated through targeted ASO binding.

## Discussion

Viral frameshifting elements are of considerable interest due to their ability to adopt diverse topologies that play a crucial role in regulating and sustaining viral proliferation. Numerous studies have explored the structural landscape of the SARS-CoV-2 FSE [8–10,36–40], including reports highlighting length-dependent structural ensembles that contribute to its functional diversity [11,41]. Because of its therapeutic importance, the SARS-CoV-2 FSE has received extensive characterisation. However, the precise FSE architectures of other viruses—such as JEV, WNV, HCV (all members of the *Flaviviridae* family), and HIV (a retrovirus) remain to be thoroughly elucidated. In this study, we initially investigated the conformational architecture of 85nt FSE from four different viruses using three independent algorithms: simRNA, trRosettaRNA, and AlphaFold3. Since, our initial analysis indicated higher similarities between the models predicted from simRNA and AlphaFold3, we used these two algorithms to call length- and ASOs- dependent conformational ensembles of viral FSEs in our further study. This approach offered a unique opportunity to perform both intra-viral and inter-viral comparative analyses of FSE structures. Additionally, we demonstrated that the native architecture of these viral FSEs is disrupted upon co-folding with various ASOs.

Although these viruses belong to distinct families, initial comparative analyses of their frameshift elements (FSEs) revealed the highest sequence homology among HIV, JEV, and WNV. In contrast, the HCV FSE showed markedly lower sequence similarity with the others, reflecting substantial per-nucleotide divergence. Interestingly, phylogenetic analysis reveals that the FSEs of WNV and JEV appear to have evolved in parallel over the course of evolution (Fig. S13). We further observed that the HCV and HIV FSE originated from one common ancestor, with the former evolving much earlier when compared to the HIV FSE (Fig. S13). Notably, while the slippery sites in most flaviviruses—including JEV, WNV, Chikungunya virus (CHKV), and Theiler’s Murine Encephalitis Virus (TMEV) [15,16,41,42] typically feature poly-U tracts (Fig. 1b, Fig. S1), both HCV and Zika virus (ZKV), despite being members of the Flaviviridae family, possess slippery sequences characterized by poly-A tracts [17,43]. Although the implications of varying slippery sequences in PRF remain unclear, all the aforementioned viruses are known to undergo -1 programmed ribosomal frameshifting (-1 PRF), except for the Zika virus, where this phenomenon is yet to be determined. Interestingly, HCV not only exhibits -1 PRF but is also reported to utilize -2/+1 and -1/+2 frameshifting events to produce two distinct proteins [17].

As stated above, simRNA, trRosettaRNA, and AlphaFold3 were simultaneously used to determine the three-dimensional (3D) architecture of all FSEs of variable lengths (85nt, 100nt (5’ and 3’), 115nt). To validate the models generated by the aforementioned software, we also modelled the 85-nt SARS-CoV-2 FSE in 3D using the same algorithms and compared them to the available cryo-EM structure (PDB ID: 6XRZ). The 3D models predicted by simRNA showed RMSD values ranging from 0 Å to 1.4 Å relative to the cryo-EM structure, indicating a high degree of similarity (Fig. 9A). Comparable trends were observed for models generated using trRosettaRNA (RMSD: 1.1–1.5 Å) and AlphaFold3 (RMSD: 0.7–1.3 Å) (Fig. 9B & 9C). Collectively, these results suggest that the models produced by all three methods are of high confidence (Fig. 9).

**Figure 9.**
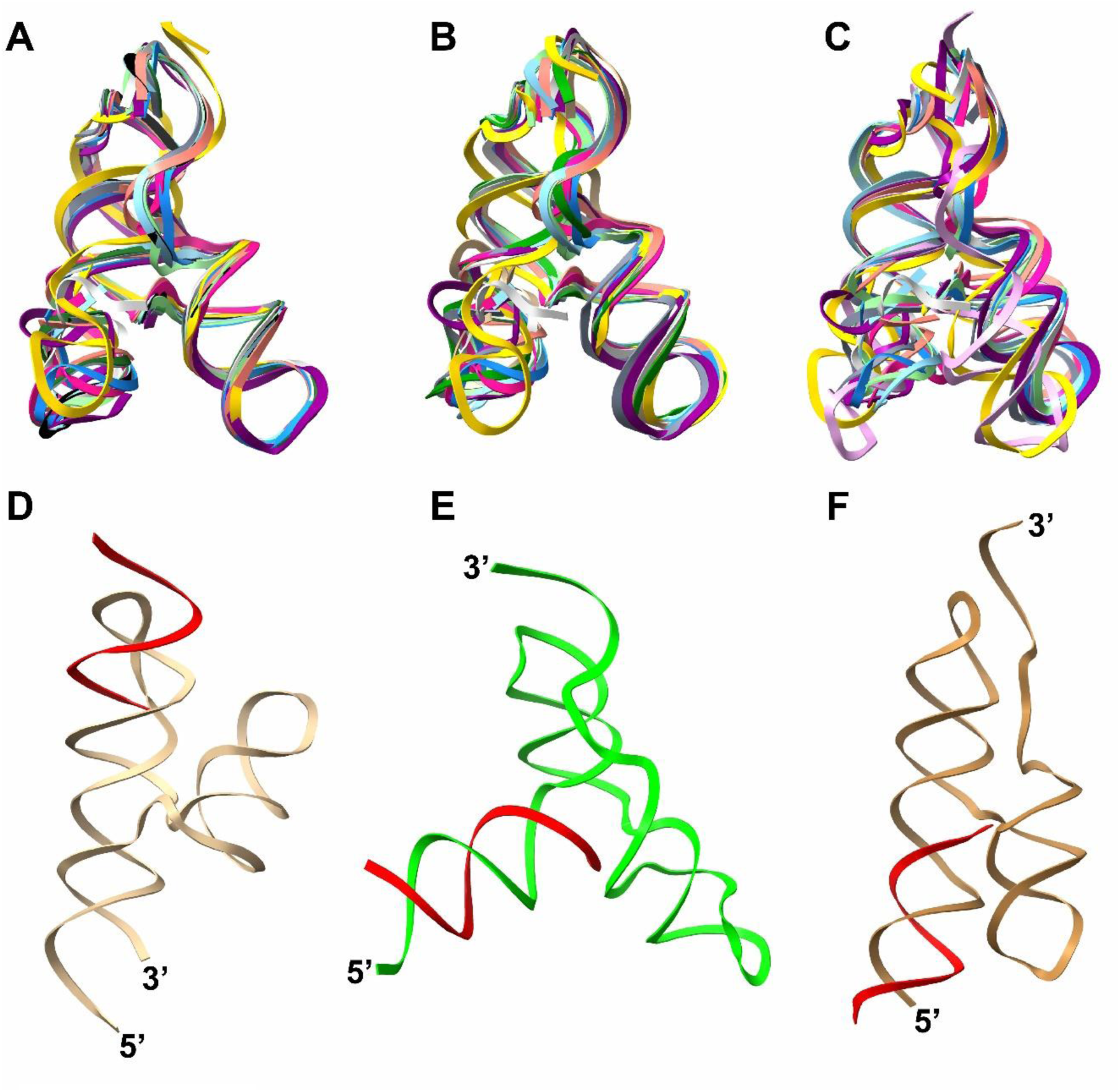
Tertiary structure determination of SARS-CoV-2 88nt FSE using **(A)** simRNA, **(B)** trRosettaRNA, and **(C)** AlphaFold3. Superimposition with the cryo-EM structure (PDB ID: 6XRZ) revealed that the computationally predicted model closely matches the experimentally derived model. This strong agreement validates the reliability of the computational algorithm used in this study for analysing other viral FSEs. Co-folding analysis of SARS-CoV-2 with already reported antisense oligonucleotides (red color) [9] **(D)** Stem 1 Disrupter-2 (S1D-2), **(E)** Slippery site 2 (Slp2), **(F)** Scrambled ASO. The binding of S1D-2 and Slp2 results in a decrease in frameshifting efficiency of SARS-CoV-2, which remains unaffected in the presence of scrambled ASO [9].

We further validated our models using molecular simulation studies, which yielded conformations closely resembling the initial predictions, as reflected by lower RMSD values. Consistent with this, comparative arc-based base-pairing analyses revealed substantial structural concordance between viral FSEs folded using simRNA and AlphaFold3.

In our study of the FSEs, we identified the presence of length-dependent conformational ensembles that emerged as nucleotides were added to either end of each FSE. Notably, our analysis revealed a conserved architecture among the core FSEs region (85 nt) across these viruses. To the best of our knowledge, this is the first report detailing the pseudoknotted architecture of the 85-nt JEV FSE. Importantly, we observed significant structural rearrangements in all viral FSEs upon adding flanking nucleotides, forming distinct, length-dependent conformational ensembles which occasionally also involve long-range tertiary interactions. Similar behaviour has previously been reported for the SARS-CoV-2 FSE using SHAPE and DMS-MaP probing [10,11]. The wild-type 85-nt SARS-CoV-2 FSE was shown to exist in equilibrium among three distinct conformations, all of which are essential for optimal frameshifting efficiency [11]. Remarkably, disruption of this equilibrium by stabilising any single conformation was found to abolish frameshifting altogether. The addition of nucleotides at either end of the FSEs led to the emergence of alternative topologies, illustrating various conformational switches regulating SARS-CoV-2 frameshifting efficiency [11,12].

Although this is the first study to characterise the length-dependent conformational ensembles of the JEV FSE, their direct impact on JEV frameshifting efficiency remains to be elucidated. However, based on prior evidence demonstrating the role of conformational switches in regulating programmed ribosomal frameshifting in the SARS-CoV-2 and WNV FSEs [11, 20], it is likely that the structural plasticity observed in the JEV FSE plays a crucial role in modulating protein translation from its frameshifting region. Our length dependent conformational analysis on JEV FSE has predominantly determined knotted conformations for the core region (85 nt) with long range interactions within 5’ and 3’ end. JEV PRF is known to produce NS1’, an alternate protein of NS1, which is expressed during the late phase infection stage of JEV [59]as opposed to NS1 expressed during the early stage of viral infection and concentrates on endoplasmic reticulum of host cells [60]. NS1’ promotes pathogen virulence by forming tunnelling nanotubes between adjacent cells [61]. These tunnels aid in the spread of viral particles and protect them from neutralizing antibodies thus enhancing virulence [61]. It seems that NS1’ expression is a tightly regulated process which is controlled by the formation of stable pseudoknots instead of least stable unknotted secondary structure. Indeed, earlier study involving surface staining analysis detected higher surface expression levels of NS1 as compared to NS1’, underlining its tightly regulated expression [62].

Previous studies on WNV FSEs of varying lengths have reported comparable structural dynamics, where the RNA adopts either a stem-loop helix or a pseudoknotted conformation depending on the number of nucleotides present [20]. Based on these observations, a model is proposed in which these two conformations act in tandem as molecular switches, regulating frameshifting and thereby modulating the expression of WNV structural and non-structural proteins at different stages of the viral replication cycle [20]. These earlier findings support our structural analysis of the WNV FSE, as we similarly observed length-dependent conformational flexibilities with knotted conformations being the dominant one. Akin to JEV, WNV -1 PRF also results in the expression of NS1’ protein, in the absence of which the virus shows reduced neuroinvasiveness and lower viral replication and RNA level [63]. Though the expression level of WNV NS1’ relative to NS1 is still obscure, given the closer proximity of WNV to JEV, it is probable that the expression of WNV NS1’ is also a tightly regulated process which is being further controlled by the stable pseudoknotted conformations of WNV FSE.

Our analysis of the HCV FSE did not reveal any pseudoknots, either in the 85-nt construct or in its extended forms. This finding aligns with recent SHAPE-MaP studies on HCV RNA, which identified the only three pseudoknots located outside the viral frameshifting region [44]. Although the structural differences between the 85-nt HCV FSE and its extended variants are subtle, and HCV FSE is known to mediate multiple types of ribosomal frameshifting (-1, +1, and +2), with direct evidence of the role of FSE conformational flexibility in this variable frameshifting phenomenon still lacking. Nonetheless, earlier studies have reported the presence of a stable stem-helix loop structure capable of stalling translating ribosomes to promote -1 PRF in HCV, while +1 frameshifting appears to depend on both the stability of the ribosome–RNA interaction and specific RNA structural features [17,21]. Presence of HCV FSE in unknotted conformation may also explain its unique pattern of PRF in HCV which is different from the other flaviviruses discussed in this study (JEV and WNV). The −2/+1 programmed ribosomal frameshifting (PRF) event in HCV leads to the production of an alternative F protein during the early stages of infection [64]. These F proteins have been shown to induce apoptosis in human dendritic cells, thereby weakening host immune responses and facilitating viral persistence [65]. In addition, F proteins contribute to immune evasion by suppressing interferon signaling, specifically through interference with RIG-I–mediated interferon induction, ultimately enhancing cellular permissiveness to infection [66].

Given that immune evasion is critical during the initial phase of HCV pathogenesis, the adoption of unknotted conformations by the HCV frameshift element (FSE) may promote efficient ribosomal frameshifting and subsequent expression of immune-modulatory proteins. Furthermore, the positioning of the HCV FSE near the 5′ end of the genome, along with the presence of a stem–loop helix, likely facilitates its efficient unwinding by translating ribosomes, ensuring proper translation of downstream viral proteins.

Recent studies using RNA-as-Graph (RAG) theory on the core HIV-1 FSE revealed that this region adopts a two-way stem-loop junction conformation. [45]. In our analysis, the 85-nt HIV FSE exhibited a similar conformation when modelled using either simRNA, trRosettaRNA and AlphaFold3, although stem II in the trRosettaRNA-derived structure appeared highly dynamic. While RAG theory did identify pseudoknots in HIV-1 FSEs, these were observed only in longer constructs. Consistently, our study also detected pseudoknot-stabilized structures in extended versions of the HIV FSE, however the core region of HIV FSE (85nt) adopted unknotted structures. -1 PRF in HIV FSE results in the expression of pol protein from the *gag* ORF. Maintaining the correct ratio of Gag-pol (20:1) protein is critical as under-/over-expression of pol protein can be detrimental to viral assembly and maturation process during the late phase of viral infection [67]. In this context, adopting a simple stem-loop helical structure by HIV FSE is plausible in fine tuning the expression of pol protein during HIV infection.

Although it remains unclear why some viral frameshifting elements (FSEs) favor highly stable pseudoknotted conformations while others adopt simpler, less stable stem–loop structures, these differences likely reflect distinct functional requirements. The greater thermodynamic stability of pseudoknots makes them more resistant to ribosomal helicase activity, thereby generating higher ribosomal tension, enhanced stalling, and increased frameshifting efficiency. However, FSEs are inherently dynamic and do not exist exclusively in a pseudoknotted state. For instance, the SARS-CoV-2 FSE has been shown to interconvert between knotted and unknotted conformations, with this structural toggling contributing to optimal frameshifting efficiency [11]. In contrast, stem–loop helices, being less stable, are more readily unwound by the ribosome, resulting in comparatively moderate frameshifting levels. This structural flexibility enables context-dependent, fine-tuned regulation of alternative protein expression, as observed in viruses such as HCV and HIV. Earlier, Lee et al. [12] proposed that a translating ribosome can remodel the SARS-CoV-2 FSE through its intrinsic helicase activity during translocation. Given that the 5′ and 3′ regions in our longer constructs are stabilized by base-pairing interactions, it is plausible that these viral FSEs similarly undergo ribosome-driven conformational transitions (Fig. 10). Consequently, the conformations observed in the longer constructs are likely transient and may become unwound during ribosomal passage, enabling the FSE to adopt alternative structural states during translation and thereby function as a molecular switch (Fig. 10).

**Figure 10.**
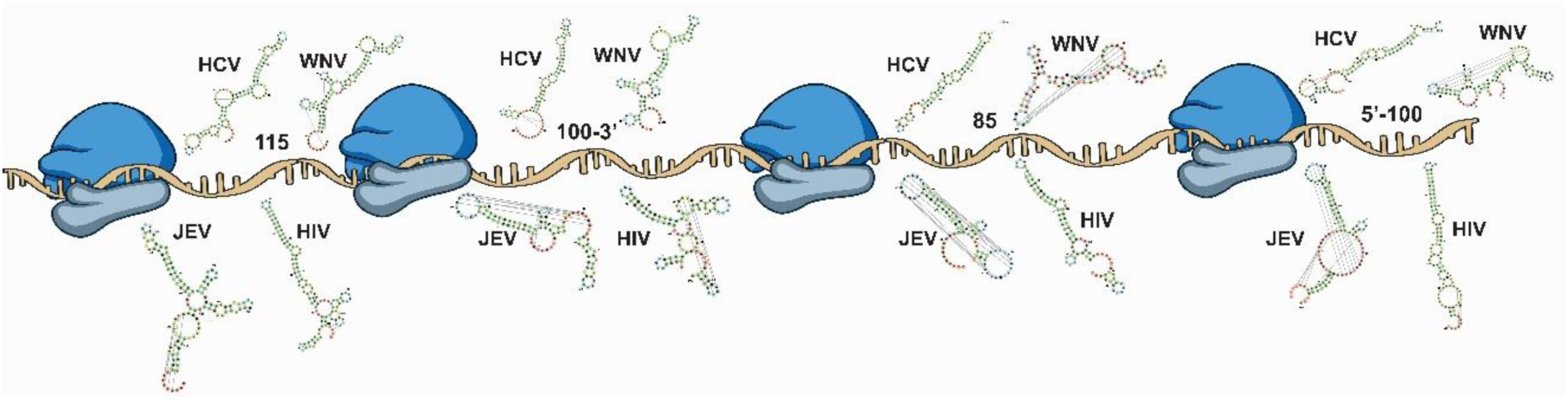
Ribosome-associated structural landscapes of viral RNA elements across genomic regions. Schematic illustration of translating ribosomes progressing along viral RNA, highlighting snapshots of local RNA structural ensembles at defined nucleotide intervals. Representative secondary structures from JEV, WNV, HCV, and HIV are depicted at ∼5′–100, ∼85, ∼100–300, and ∼115 nucleotide regions relative to the coding sequence, illustrating diverse folding architectures. The conformational variability observed in longer RNA constructs appears transient and is likely resolved during ribosomal transit, allowing FSEs to dynamically adopt alternative structures during translation and act as regulatory molecular switches.

Our PCA analysis also reveals structural heterogeneity across 85nt and 115nt generated models. simRNA derived models consistently shows, broader and more continuous conformational landscape, indicating higher structural plasticity. In contrast, AlphaFold3-initialized structures favor more confined or discrete clusters, suggesting stabilization in defined conformational states with comparatively restricted dynamics across viral FSEs. Nevertheless, both modelling approaches produced largely similar architectures, with only subtle variations observed across the viral FSEs.

The observed structural diversity prompted us to design multiple structure-disrupting ASOs, which we evaluated through co-folding analyses with their corresponding FSE targets using simRNA and AlphaFold3. These co-folding studies revealed significant structural alterations in the native FSE upon the introduction of ASO at two distinct target sites within the same FSE (either 85nt or 115nt in length).

The partition function analysis highlights that ASO binding can effectively outcompete the native tertiary folds of the viral FSE. The formation of a highly stable ASO–FSE heteroduplex (Region II), supported by dense interaction patterns and favorable negative free energy values, suggests that ASO binding is thermodynamically preferred over the intrinsic intramolecular interactions of the FSE. Concurrently, the disruption and diffusion of contacts within Region I indicate that the native tertiary architecture becomes destabilized and shifts toward a more heterogeneous conformational ensemble upon ASO engagement. Together, these observations support a model in which ASO binding overrides native folding constraints, thereby remodeling the structural landscape of the FSE.

Such structural disruption of the native FSE by ASOs could modulate frameshifting efficiency, thereby impacting viral replication and pathogenesis. In our study, co-folding of ASOs targeting the Stem-2 and slippery site of the SARS-CoV-2 FSE resulted in clear native structure disruption (Fig. 9C and 9D). In contrast, no structural changes were observed with scrambled ASOs (Fig. 9E, Fig. S14). Supporting this, Zhang et al. [9] demonstrated that these ASOs significantly reduces the frameshifting efficiency of SARS-CoV-2 FSEs and exhibited potent antiviral activity *in vitro*. Given the ability of our ASOs to perturb FSE conformations, future biochemical studies could evaluate their potential to suppress frameshifting and reduce viral replication, offering promising avenues for antiviral intervention.

While this study delineates robust length- and ASOs-dependent conformational landscape of viral frameshifting elements using complementary computational approaches, these structures represent equilibrium folds in the absence of active translation and cellular cofactors. Future integration of ribosome-coupled structural probing and in-cell validation will be essential to directly correlate specific FSE topologies with frameshifting efficiencies and viral fitness under physiological conditions.

Though this study employs multiple modeling approaches, the agreement and divergence observed between simRNA and AlphaFold3 predictions can be better understood in the context of their distinct algorithmic frameworks and inherent limitations. simRNA utilizes a coarse-grained representation with knowledge-based potentials and Monte Carlo sampling, favoring thermodynamically stable conformations and enabling explicit exploration of complex features such as pseudoknots. In contrast, AlphaFold3 relies on deep learning–based inference guided by structural priors embedded within its training data, which can bias predictions toward known structural patterns and may impose constraints on representing highly dynamic or non-canonical RNA topologies. As a result, convergence is typically observed in regions with well-defined secondary structures, whereas divergence arises in flexible regions and in the treatment of higher-order interactions, including pseudoknots. We acknowledge that each approach carries specific limitations, including differences in conformational sampling, treatment of long-range interactions, and dependence on prior structural information. To strengthen model selection and confidence, we incorporated multiple evaluation metrics, including RMSD-based clustering of simulated conformations and confidence estimates such as predicted alignment error (PAE) where applicable. Representative models were selected based on convergence within low-RMSD clusters and consistency across methods. Importantly, these computational assessments were complemented by molecular simulation studies and arc-based base-pairing analyses, which enabled systematic mapping of base-pairing interactions and facilitated the identification of structurally consistent models. Collectively, this integrative strategy improves confidence in the selected conformations while also highlighting regions of structural variability that may require further experimental validation.

The conformational plasticity uncovered in this study suggests that viral frameshifting elements function as dynamic RNA ensembles rather than static structural motifs, with sequence context and antisense binding acting as tunable regulators of its architecture. Integrating high-resolution experimental approaches, such as in-virion chemical mapping and structure probing with quantitative frameshifting assays, will be critical to establish direct structure–function relationships for individual conformers. Such efforts will not only clarify how specific FSE architectures are selected during translation but also enable rational design of structure-selective antisense modulators, positioning viral FSEs as tractable RNA targets for next-generation antiviral strategies.

## Materials and Methods

### Virus FSE sequence

To determine the conformational landscape of FSE in 3 flaviviruses and 1 retrovirus, FSE regions were retrieved from the NCBI nucleotide database. For the JEV reference sequence (Accession ID: NC_001437.1), the 85nt FSE occupies from nucleotides 3554-3638. 85nt FSE of WNV (Accession ID: NC_001563.2) occupies from 3528-3612. In HCV strain (AF011751.1), 85nt FSE spans from 354-438. In the HIV strain (NC_001802.1), 85nt FSE starts from the nucleotide position 1626 and ends at 1710. In addition to 85nt FSE regions, constructs (5’-100 {15 additional nucleotides at 5’ of FSE}, 100-3’ {15 additional nucleotides at 3’ end of FSE}, and 115 {15 additional nucleotides both at 5’ and 3’}) were also used to determine the conformational flexibility of FSE regions (Fig. 1A) across various algorithms and viruses.

### IPknot for FSE secondary structure predictions

To determine the secondary structure, especially pseudoknots, if any are present in FSE regions of JEV, WNV, HCV, and HIV, we used the web server version of IPknot [46]. The RNA sequence was provided in FASTA format, and Level 2 (nested pseudoknots) with the McCaskill model was used for secondary structure prediction. IPknot output for each construct used in this study is shown in Table S1 and also available through zenodo repository: https://doi.org/10.5281/zenodo.17432064 in dot-bracket notation form.

### FSE tertiary structure predictions and simulations

#### simRNA

To determine the three-dimensional architecture of FSE of various viruses used in this study, we performed RNA simulation studies using simRNAweb 2.0 [32]. simRNA utilises a coarse-grained model and the Monte Carlo method to statistically determine energies to determine the tertiary structure of a given RNA sequence accurately. simRNA was run in default mode after providing the fasta sequences of each FSE and their respective IPknot output (.dbn file) as soft secondary structure restraints to perform simulations for tertiary structure generation (Table S1 and zenodo repository: https://doi.org/10.5281/zenodo.17432064).

To understand the effect of ASOs on the predicted tertiary models of FSE, we ran co-modelling simulations using the two-chain RNA option in simRNA. Each FSE construct was co-modelled with its respective ASO (identified and clustered through OligoWalk, see below) using hard secondary structure restraints. These rigid secondary structure restraints were obtained after running both the FSE sequence (FASTA sequences) and its respective ASO sequence through the RNAcofold webserver of ViennaRNA web services [47].

#### trRosettaRNA

We also predicted the tertiary structure of our FSE using trRosettaRNA [33]. trRosettaRNA uses an automated deep learning approach to predict the tertiary structure of RNA using energy minimisation. Once the RNA sequence (as a FASTA file available through the Zenodo repository (https://doi.org/10.5281/zenodo.17432064) is provided, trRosettaRNA calculates the inter-nucleotide distance distributions, which are further converted into smooth restraints for 3D structure prediction based on energy minimisation. For most FSE constructs, the trRosettaRNA predicted three-dimensional (3D) model with medium confidence.

#### Alphafold3

We also used Alphafold3 [34] simulations for the predictions and comparative analysis of the 3D models of our FSE constructs. Alphafold3 works in tandem with two modules: 1) Pairformer, and 2) Diffusion Module. The Pairformer module determines the paired and single relationship between the provided sequences by a conditioning network. Pair representation identifies molecular relationships; single representation focuses on individual molecular components. This collective information is passed onto the Diffusion module for model building. The diffusion module uses raw atomic coordinates to predict the final accurate model after being conditioned using the single and pair representations obtained from the pairformer module. All jobs were run in default mode by providing one copy of each RNA sequence (available through zenodo repository https://doi.org/10.5281/zenodo.17432064. For the co-folding analysis, we used RNA sequences of individual FSE constructs and their corresponding ASOs, identified through OligoWalk (see below), in AlphaFold3.

### Molecular Dynamics Simulations

All-atom molecular dynamics (MD) simulations were performed using GROMACS 2025.4 [51] with the AMBER 94 force field [52,53], optimized for RNA structures. Tertiary models of viral RNA FSE constructs for JEV, WNV, HCV, and HIV (85 nt and 115 nt) generated using simRNA and AlphaFold3 were used as inputs for each simulation study. Prior to the run, each model system was solvated in a cubic box using the TIP3P water model [54] with a minimum distance of 1.0 nm from the box edge. Sodium and chloride ions were added to neutralize the system and achieve a 150 mM physiological ionic strength.

Energy minimization was performed using the steepest descent algorithm (maximum force < 1000 kJ/mol/nm). The system was equilibrated in two phases: NVT (constant volume) for 100 ps at 300 K using the V-rescale thermostat [55], followed by NPT (constant pressure) for 100 ps at 1 bar using the Parrinello-Rahman barostat [56]. Production MD simulations were conducted for 100 ns in the NPT ensemble with a 2 femtosecond (fs) timestep. Long-range electrostatics were calculated using the Particle Mesh Ewald (PME) method [57], with van der Waals interactions truncated at 1.0 nm. LINCS algorithm [58] constrained all bonds involving hydrogen atoms. Coordinates were saved every 10 ps for subsequent analysis.

### Principal component analysis (PCA) analysis

PCA was used to identify ensembles arising from the backbone motions in the 85-nt and 115-nt simulated RNA structures derived from simRNA- and AlphaFold3. For each trajectory, frames were processed using MDAnalysis (Universe-based trajectory handling) [70,71] and phosphate backbone atoms were selected as the structural descriptor. MD frames were subsampled at a fixed stride (STRIDE = 5) to reduce temporal correlation and to match the point density used for plotting. Each retained frame was least-squares superposed onto the 0 ns reference structure (frame 0) by applying a Kabsch rigid-body alignment (singular value decomposition [SVD]-based rotation) to the selected phosphate coordinates, thereby removing overall translation and rotation prior to PCA [72]. The aligned coordinate matrix (frames × 3N Cartesian coordinates) was mean-centered and decomposed by singular value decomposition (SVD), which is equivalent to diagonalizing the covariance matrix of positional fluctuations; the resulting principal components (eigenvectors) and associated variances (eigenvalues) were used to compute per-frame PC scores and the percentage of total variance explained by each component, following standard essential-dynamics methodology. Trajectory frames were projected onto PC1–PC2 to visualize the dominant modes of backbone motion, and conformational sampling was stratified into three time windows corresponding to early phase (0–1 ns), intermediate phase (1–99 ns), and terminal phase (99–100 ns), with simulation time obtained from trajectory timestamps and converted from ps to ns. PCA theory for MD (“essential dynamics”) follows the canonical framework described by Amadei et al. [68], while implementation and trajectory handling follow the MDAnalysis software description [69]. This analysis follows contemporary best-practice implementations of PCA/essential dynamics for molecular simulation trajectories and employs widely used open-source tooling for trajectory handling and interpretation.

### ASO identification through OligoWalk

To identify and design ASO that can bind to our FSE constructs, we used OligoWalk, an online ASO-generating tool [48]. OligoWalk calculates the thermodynamic features of sense-antisense hybridisation by predicting changes in free energy when a ASO binds to its target RNA using the partition function [48]. The tool was run in default mode, using the target RNA sequences, except that the oligomer length was set to 21 nucleotides. ASOs identified for different FSE constructs are provided in Table S3.

## Acknowledgments

The authors would like to acknowledge Department of Biotechnology, NIPER-Raebareli, for its invaluable support, which greatly contributed to the completion of this article. KA acknowledges the Department of Pharmaceuticals, Ministry of Chemicals and Fertilisers, Govt. of India, for providing fellowship assistance. AD acknowledges the Department of Biotechnology, Govt. of India, for the Ramalingaswami Re-entry Fellowship (BT/RLF/Re-entry/02/2021) and ANRF for Early Career Research Grant (ECRG) (ANRF/ECRG/2024/001002/LS). This article bears NIPER-R communication number 768.

## Data Availability statement

FASTA (.fa), clustal (.aln) and dot-bracket notation (.dbn) which were used as an input files to run simRNA, trRosettaRNA and Alphafold3 are accessible through Zenodo repository (https://doi.org/10.5281/zenodo.17432064). Code for generating heatmaps usings R programming are also available through the Zenodo repository (https://doi.org/10.5281/zenodo.17432064). Predicted Aligned Error (PAE) files of AlphaFold3 in, JSON format is also available through Zenodo repository (https://doi.org/10.5281/zenodo.17432064). Atomic coordinates of the frameshifting models reported in this study, are accessible through Zenodo repository (http://DOI 10.5281/zenodo.17261773).

