## Supplementary File for "Pan-Viral Conformational Landscapes of Frameshifting Elements Reveal Length-Dependent Plasticity and Antisense-Driven Structural Reprogramming"

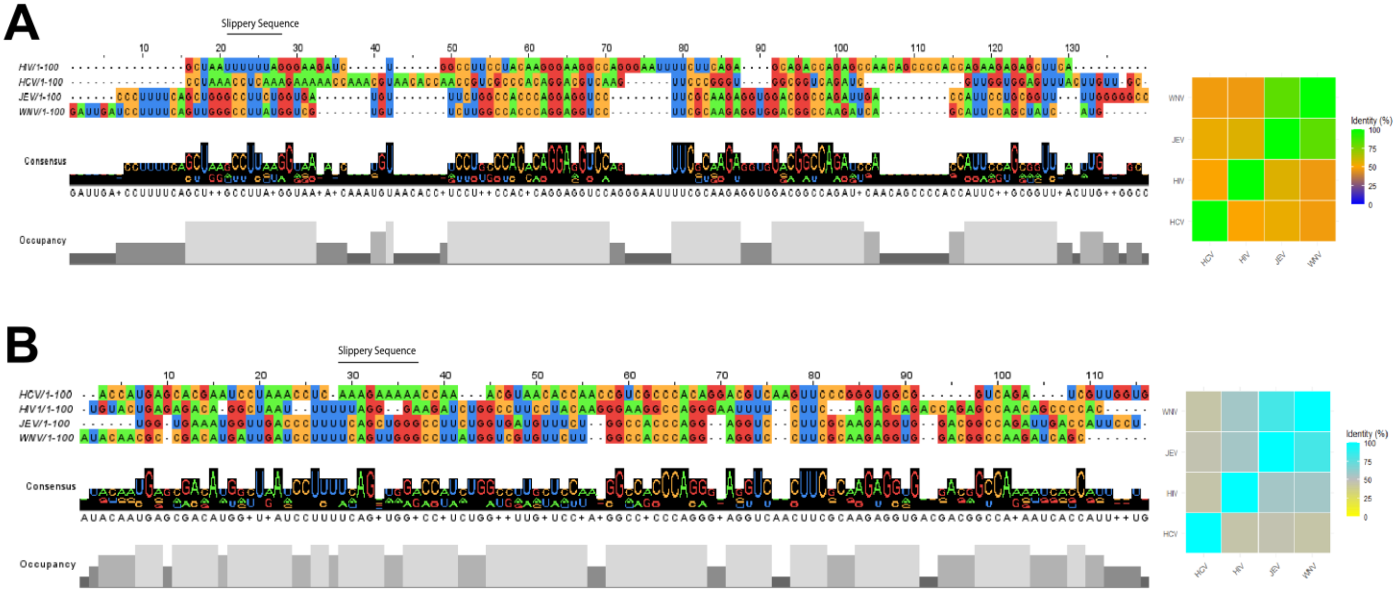


**Figure S1:** Sequence alignment and per cent identity matrix in the form of a heatmap for (**A)** 100-3'nt FSE constructs, and (**B)** 5'-100nt FSE constructs of JEV, WNV, HCV and HIV. Like 85nt and 115nt FSEs, sequence alignment shows higher similarity between the FSEs (100-3' and 5'-100 constructs) of JEV, WNV and HIV.

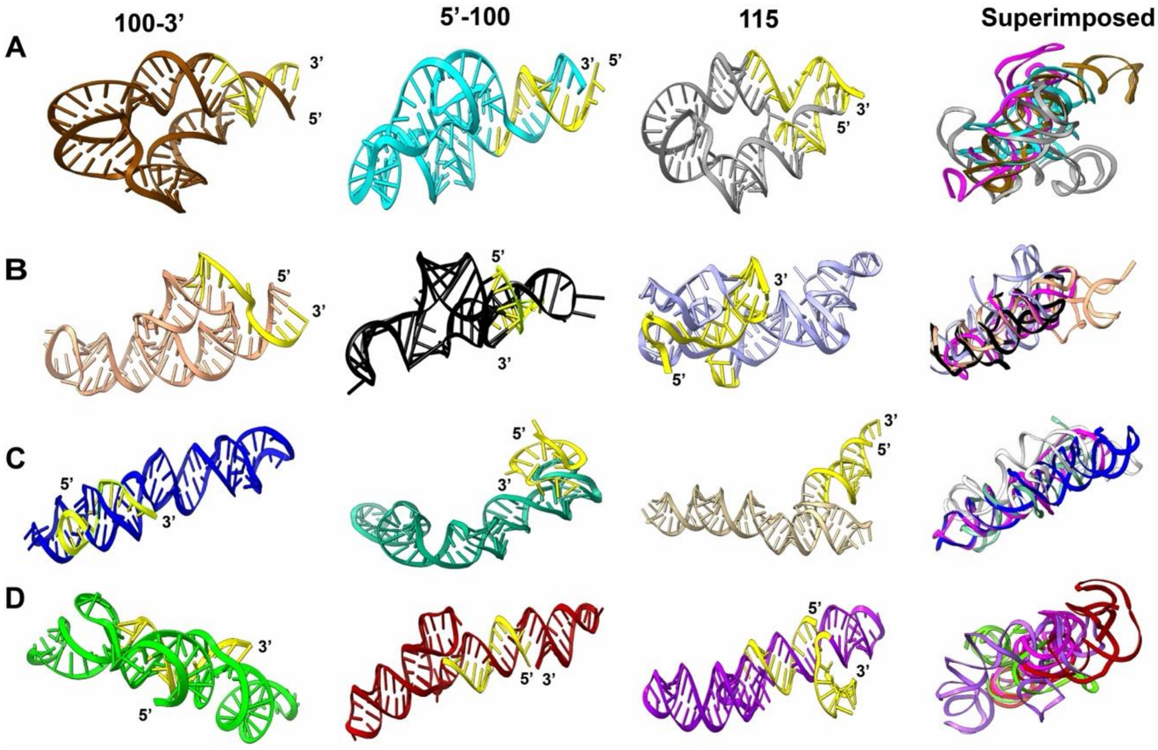


**Figure S2:** Tertiary structure models of larger FSE constructs (100-3’nt, 5’-100nt, and 115nt) for **(A)** JEV, **(B)** WNV, **(C)** HCV, **and (D)** HIV generated through AlphaFold3. The last column represents the superimposed conformations of longer FSE constructs on the 85 nt FSE region of their respective viruses generated through AlphaFold3.

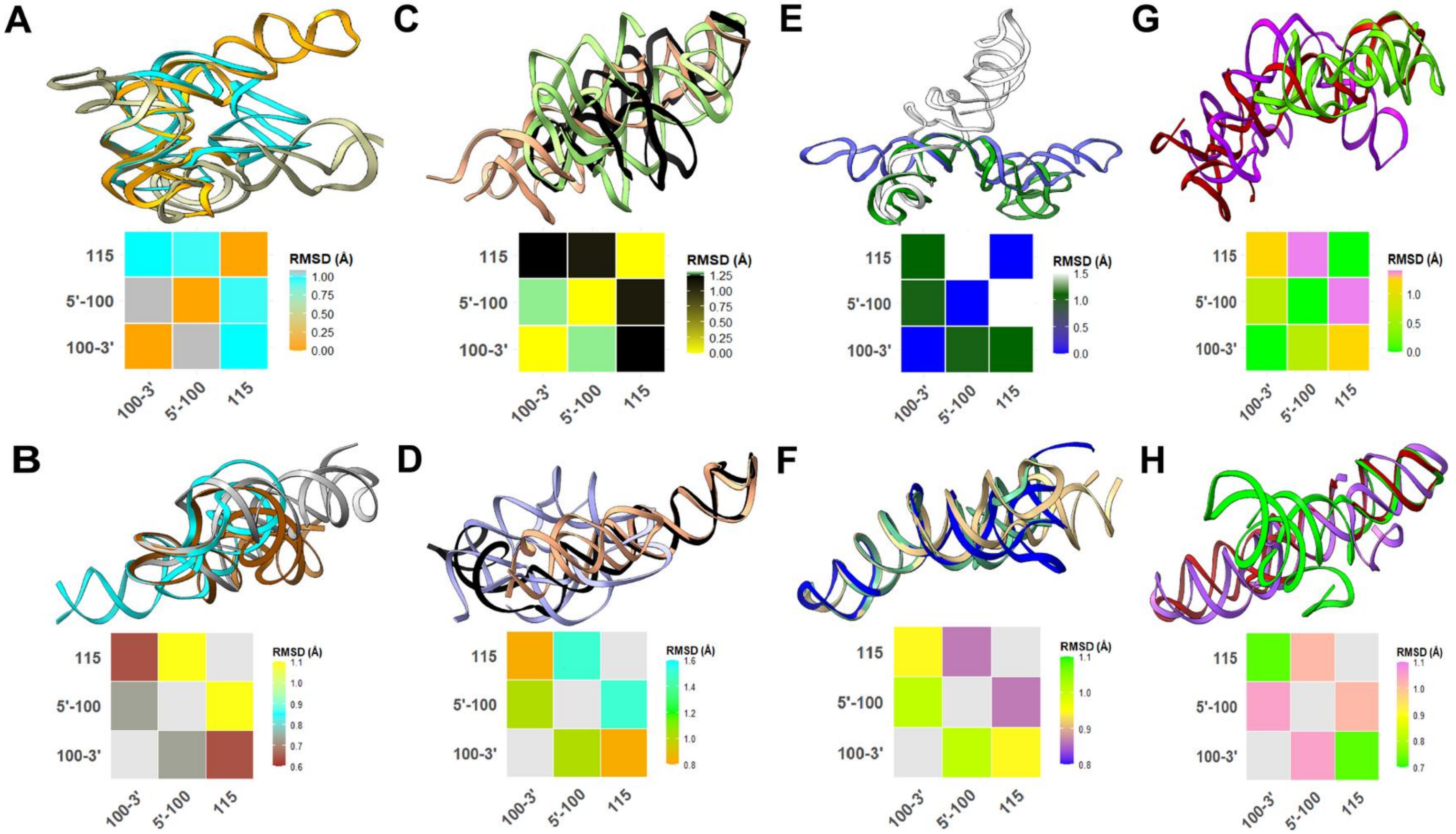


**Figure S3:** Superimposition of longer FSE constructs (100-3'nt, 5'-100nt, and 115 nt) of viral FSE generated with RMSD being represented as a heatmap, generated for JEV through (**A)** simRNA, and **(B)** AlphaFold; generated for WNV through **(C)** simRNA, and **(D)** AlphaFold3; generated for HCV through **(E)** simRNA, and (**F)** AlphaFold3; and generated for HIV through **(G)** simRNA, and **(H)** AlphaFold3.

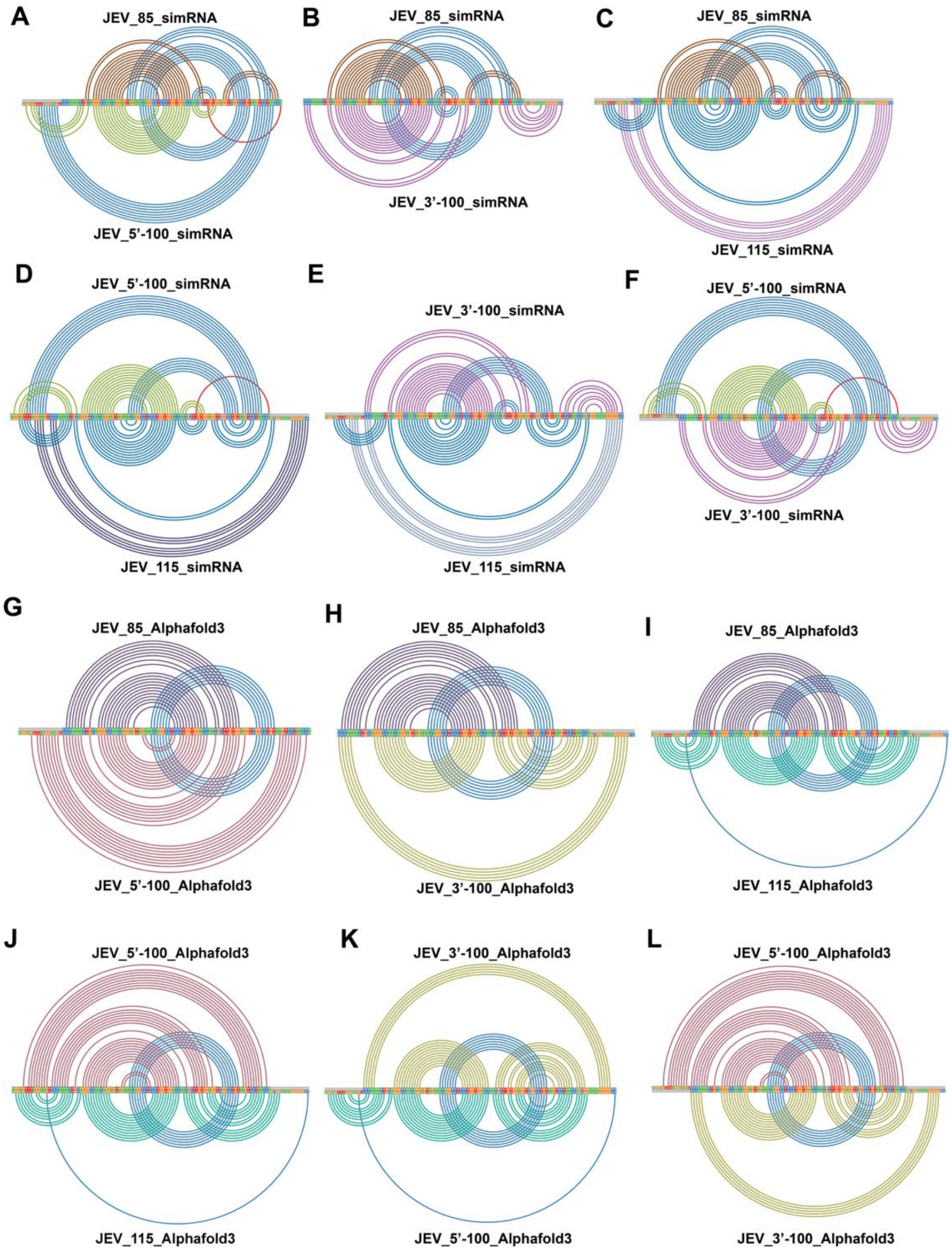


**Figure S4**: Comparative arc plot analysis between the longer constructs (5'-100, 100-3' and 115 nt) of JEV FSE models generated using **(A), (B), (C), (D), (E),** and **(F)** simRNA and **(G), (H), (I), (J), (K),** and **(L)** using AlphaFold3.

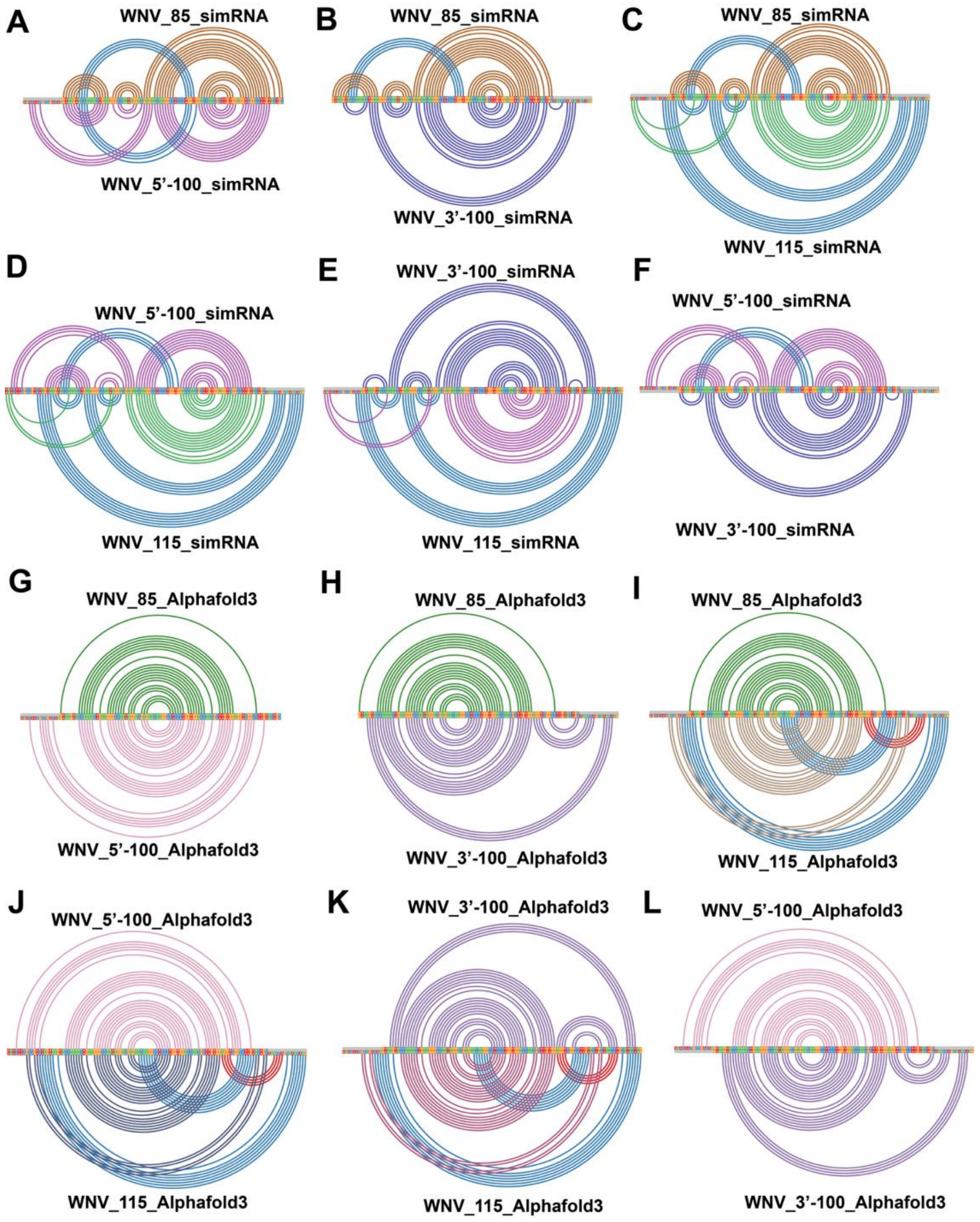


**Figure S5:** Comparative arc plot analysis between the longer constructs (5'-100, 100-3' and 115 nt) of WNV FSE models generated using **(A), (B), (C), (D), (E),** and **(F)** simRNA and **(G), (H), (I), (J), (K),** and **(L)** using AlphaFold3.

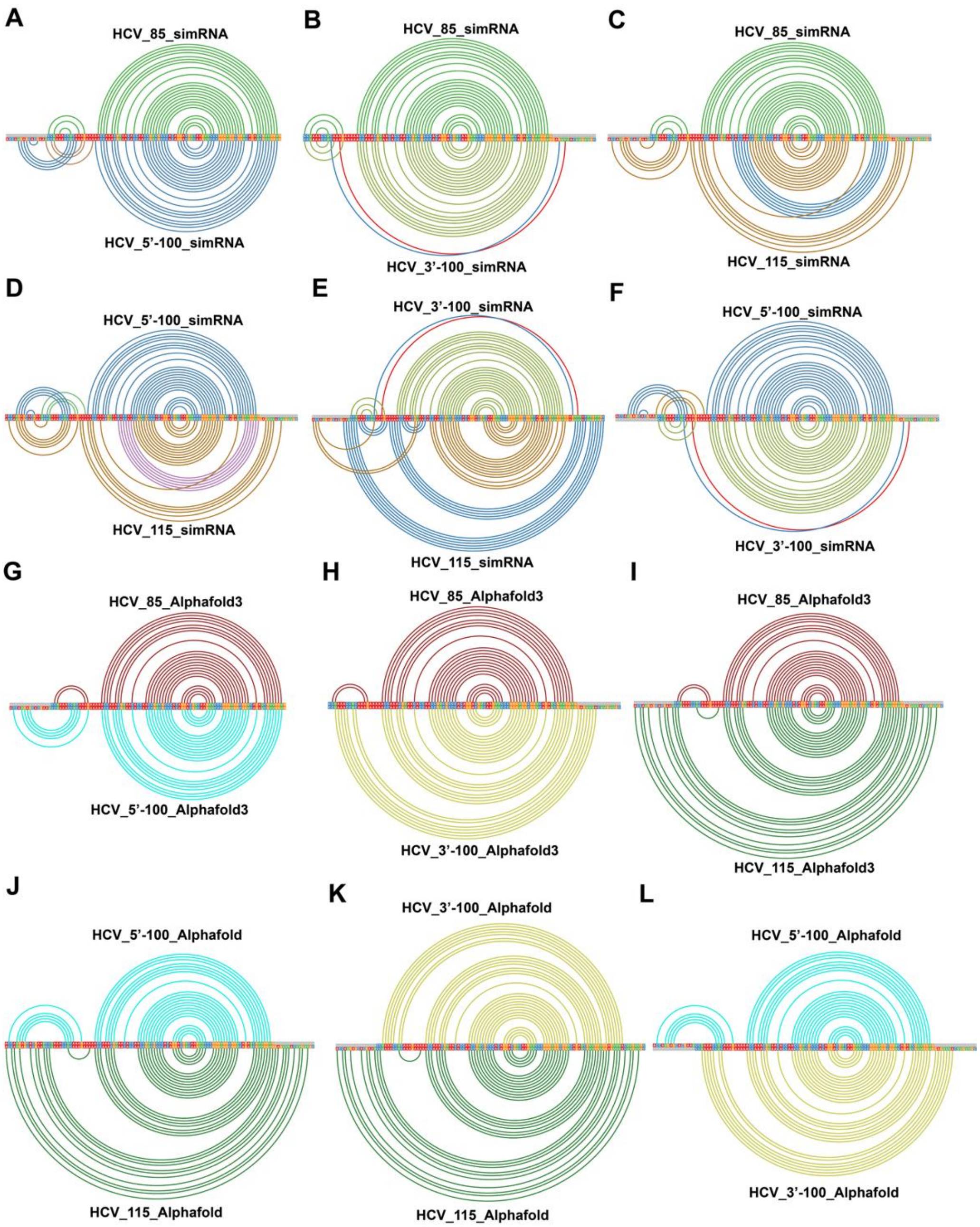


**Figure S6:** Comparative arc plot analysis between the longer constructs (5'-100, 100-3' and 115 nt) of HCV FSE models generated using **(A), (B), (C), (D), (E),** and **(F)** simRNA and **(G), (H), (I), (J), (K),** and **(L)** using AlphaFold3.

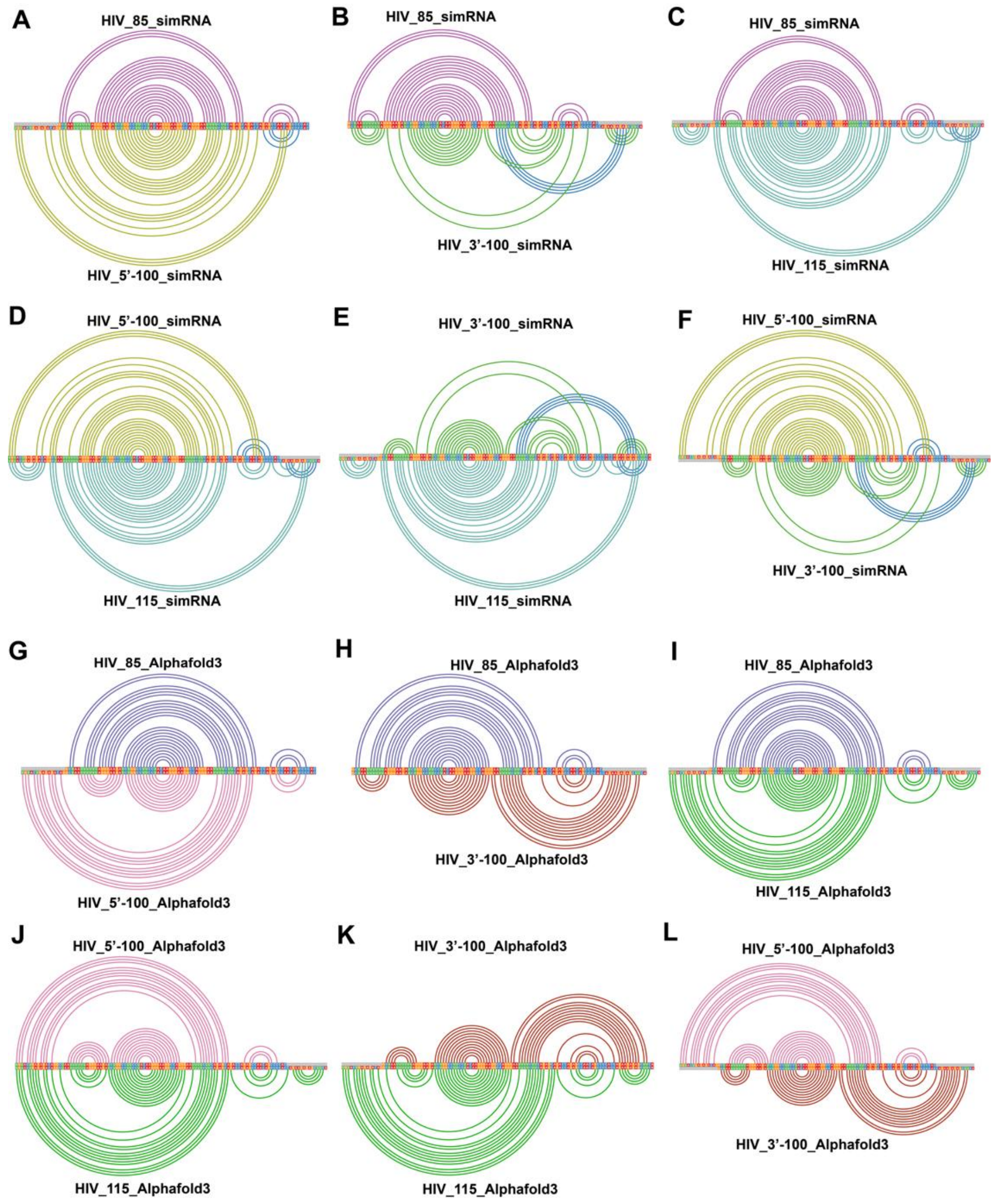


**Figure S7:** Comparative arc plot analysis between the longer constructs (5'-100, 100-3' and 115 nt) of HIV FSE models generated using **(A), (B), (C), (D), (E),** and **(F)** simRNA and **(G), (H), (I), (J), (K),** and **(L)** using AlphaFold3.

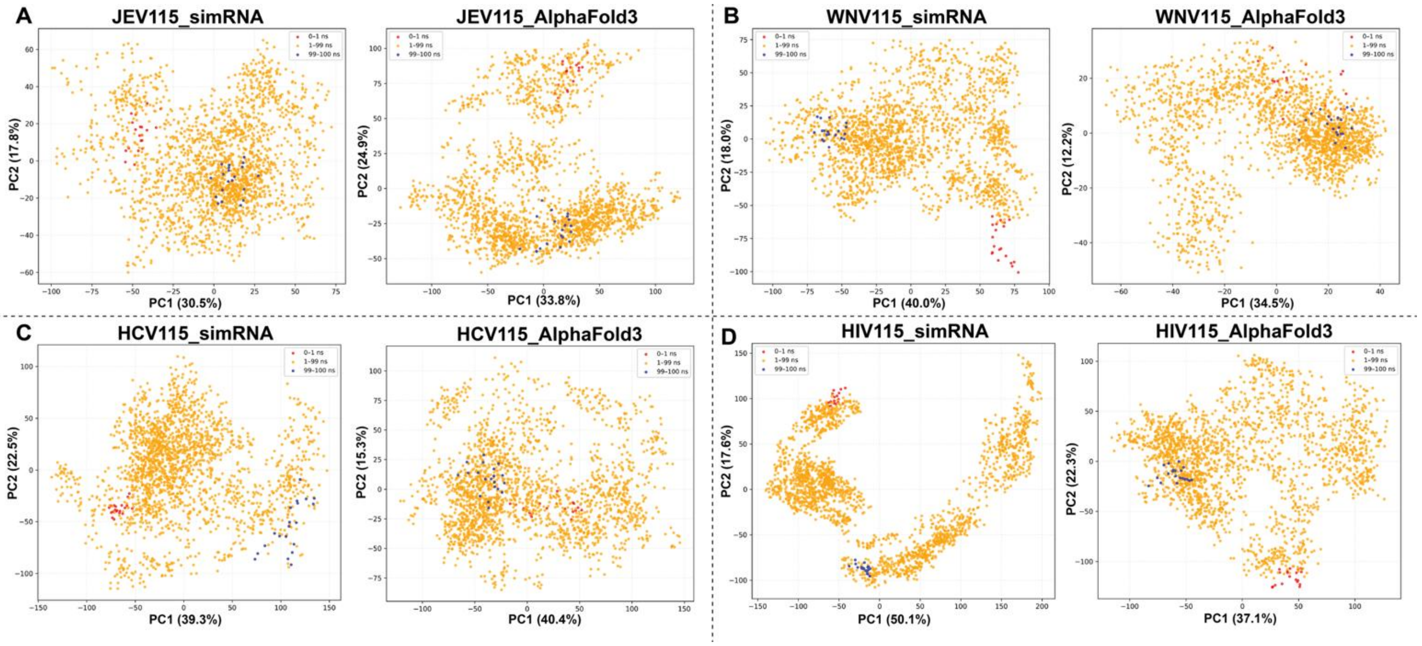


**Figure S8.** Time-resolved PCA reveals conformational ensembles of 115-nt viral FSEs modeled using simRNA and AlphaFold3 for **(A)** JEV, **(B)** WNV, **(C)** HCV, and **(D)** HIV. simRNA structures display broader dispersion and multiple basins, indicating higher heterogeneity, whereas AlphaFold3 models show tighter clustering, suggesting restricted sampling and stabilization in defined conformational states.

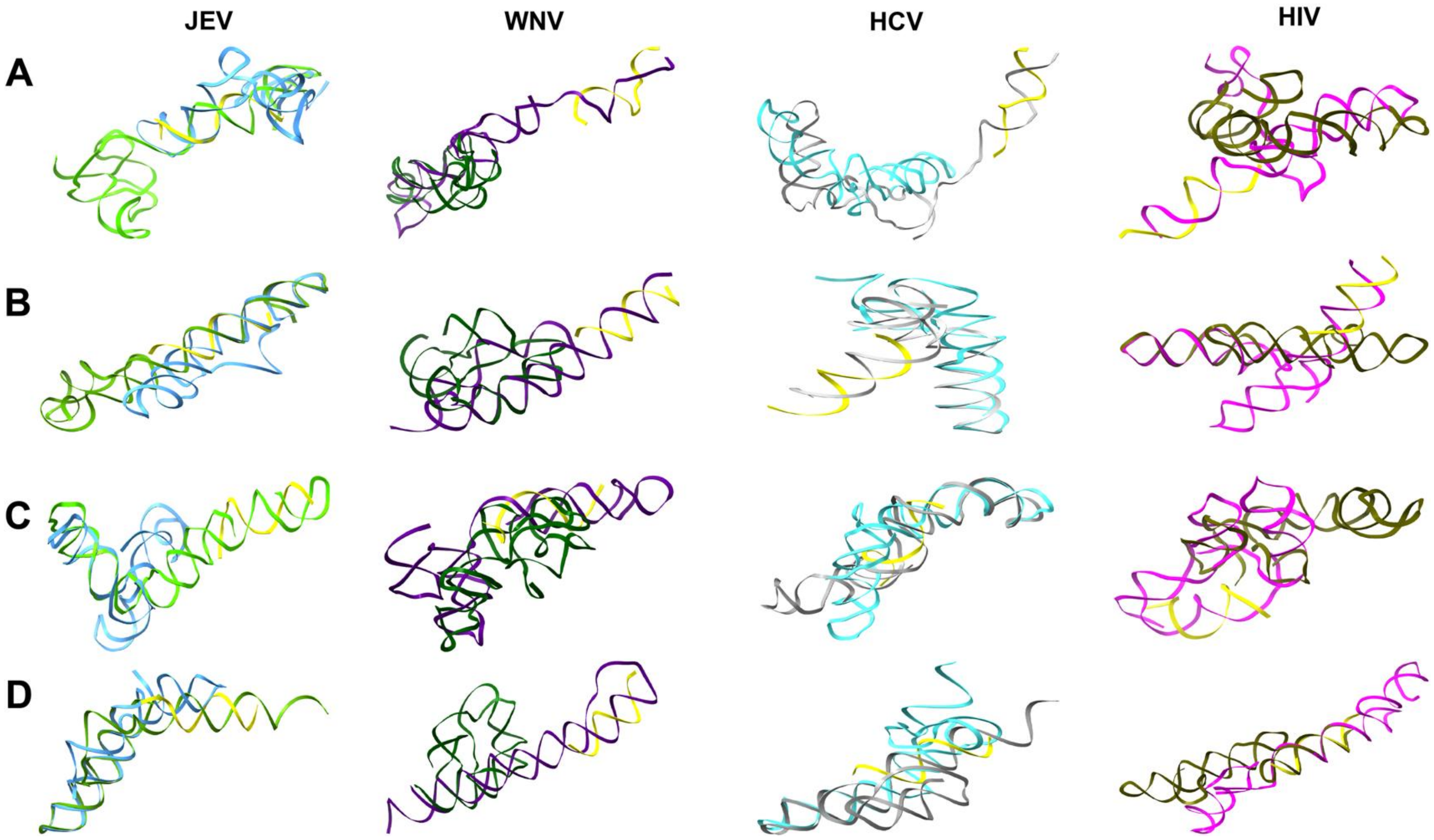


**Figure S9:** Co-folding analysis between ASO (yellow) generated through OligoWalk and binding to the 5' end of 115nt FSEs of JEV, WNV, HCV, and HIV using **(A)** simRNA and **(B)** AlphaFold3. Co-folding analysis between ASO (yellow) generated through OligoWalk and binding to the middle or 3' end of 115nt FSEs of JEV, WNV, HCV, and HIV using **(C)** simRNA, **(D)** AlphaFold3. Superimposition over native 115nt FSE shows a significant structural alteration in viral FSE in the presence of ASO.

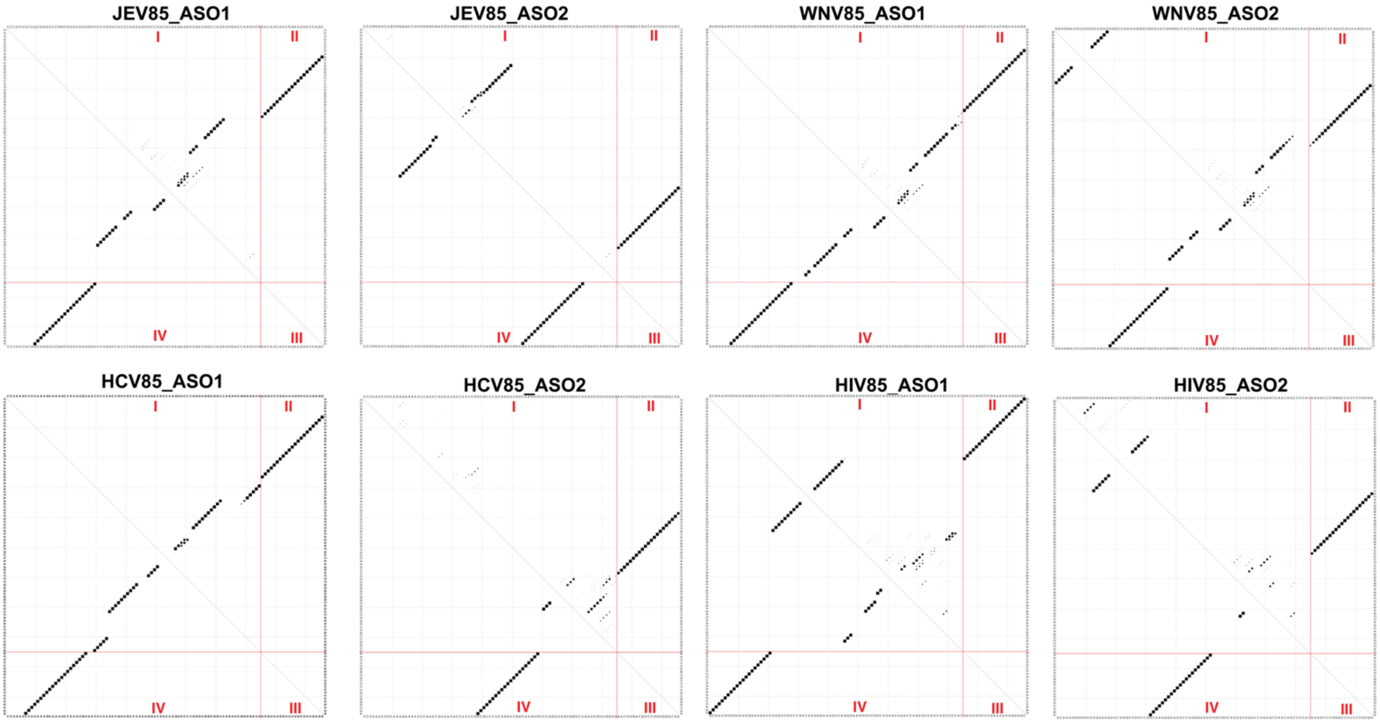


**Figure S10.** Dot-plot graphs illustrate Boltzmann-weighted secondary structures from partition function and minimum free energy analyses of viral FSEs (85 nt) co-folded with ASOs. Regions (I-IV) distinguish FSE–FSE and FSE–ASO interactions. Black blocks indicate base pairing intensity. Diffuse patterns reflect structural heterogeneity, while dense, well-defined blocks in Regions II indicate highly stable FSE–ASO interactions.

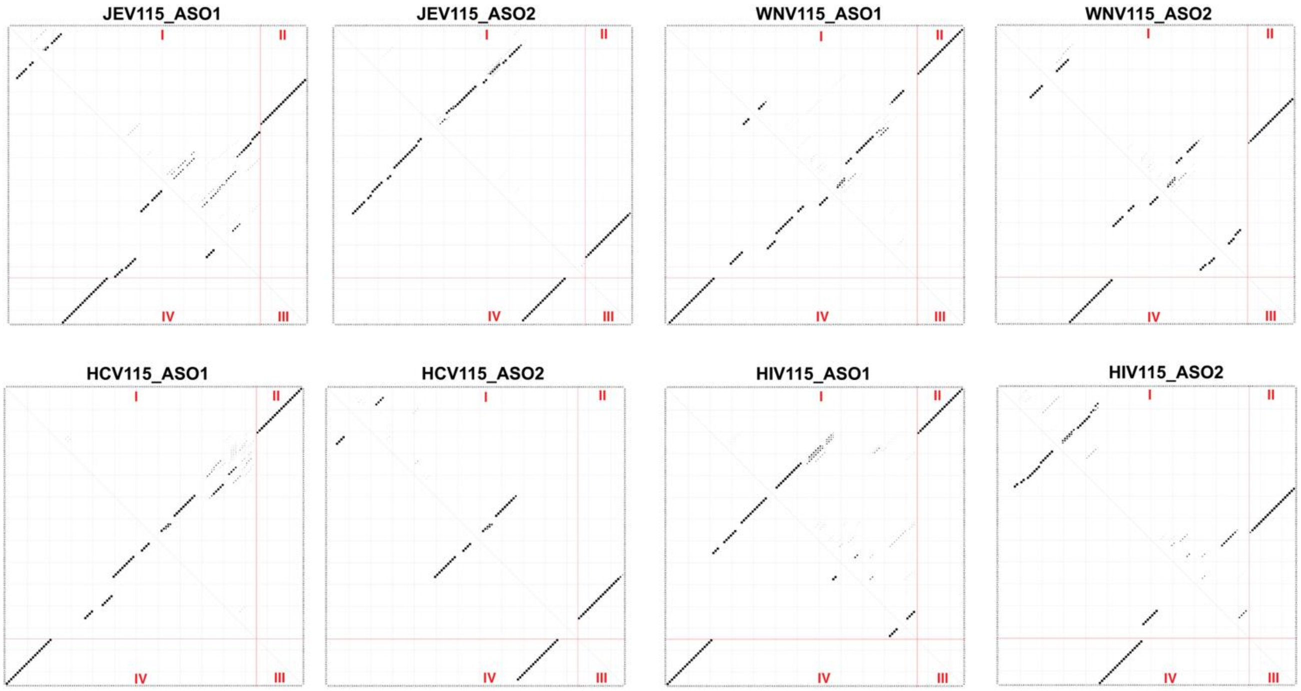


**Fig. S11.** Dot-plot graphs illustrate Boltzmann-weighted secondary structures from partition function and minimum free energy analyses of viral FSEs (115 nt) co-folded with ASOs. Regions (I-IV) distinguish FSE–FSE and FSE–ASO interactions. Black blocks indicate base pairing intensity. Diffuse patterns reflect structural heterogeneity, while dense, well-defined blocks in Regions II indicate highly stable FSE–ASO interactions.

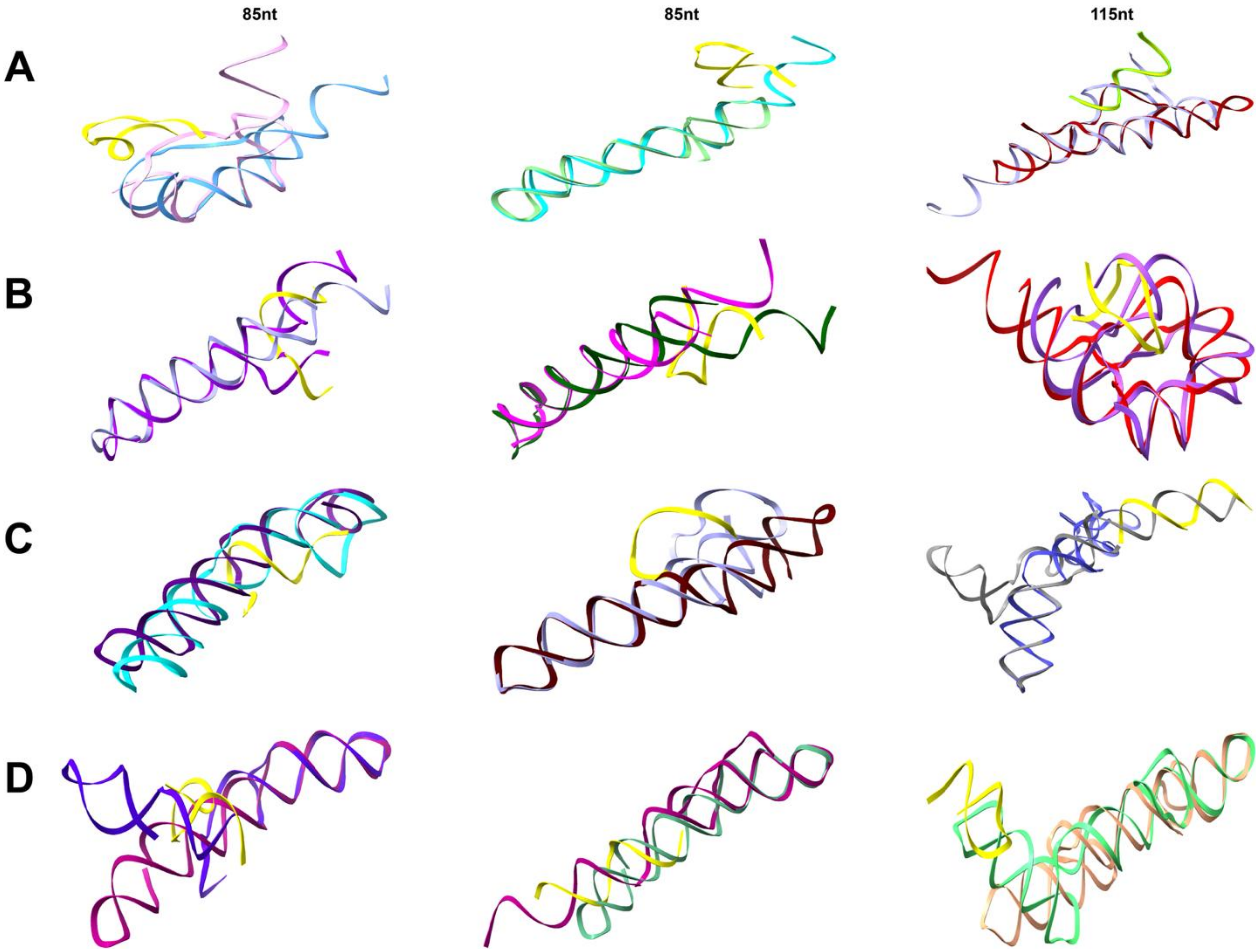


**Figure S12:** Co-folding analysis between scrambled ASOs and 85nt and 115nt FSEs of **A**. JEV, **B**. WNV, **C**. HCV and **D**. HIV generated using AlphaFold3. Superimposition with native FSE architectures shows no significant changes in their topologies, confirming the validity of our structural analysis of FSEs in the presence of various ASOs.

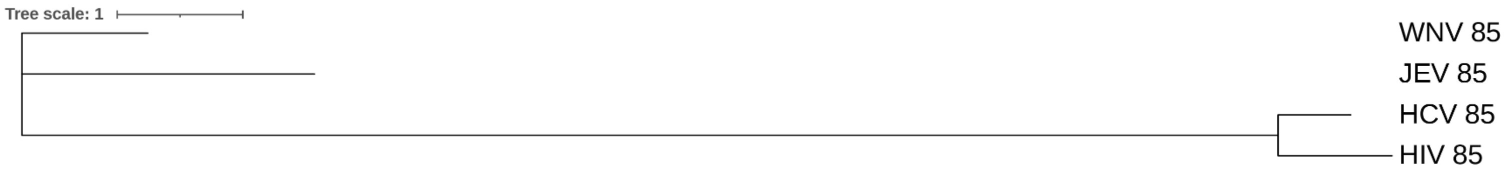


**Figure S13:** The phylogenetic tree of the 85-nt FSE sequences from JEV, WNV, HCV, and HIV suggests that JEV and WNV FSEs share a common ancestor, with JEV FSE diverging earlier. The longer branch length observed for HIV indicates that its FSE evolved later than those of JEV, WNV, and HCV during evolution.

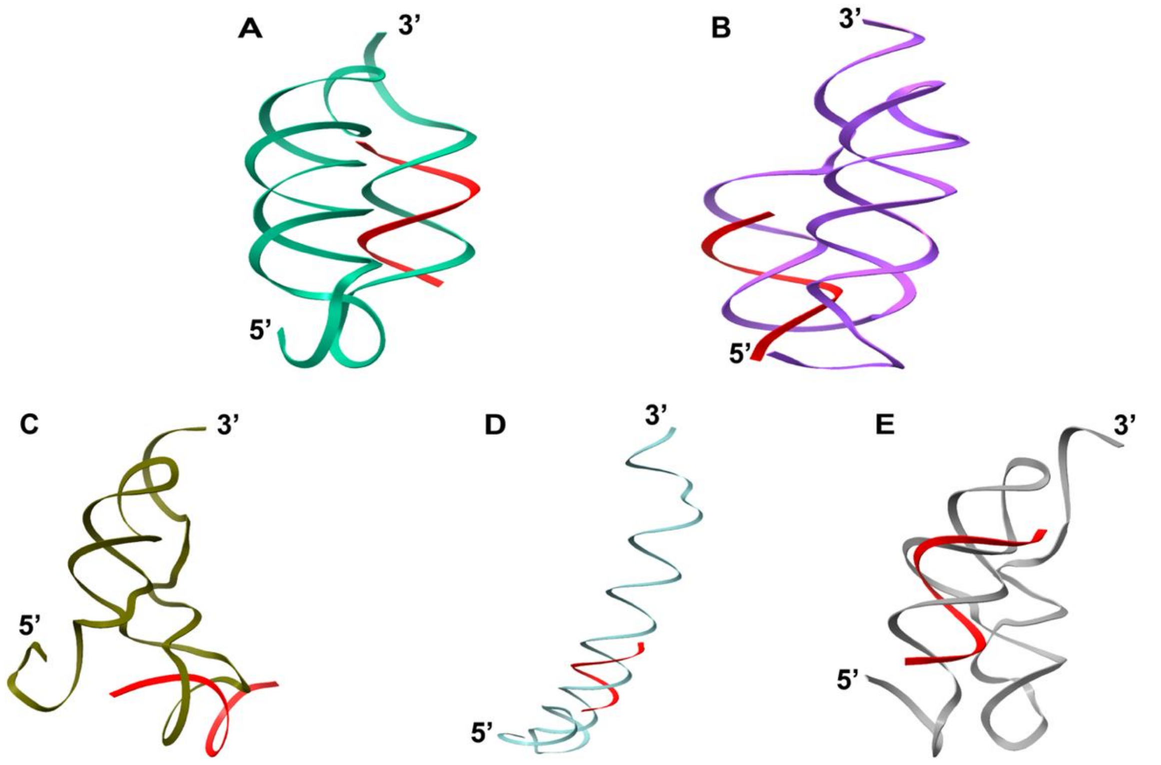


**Figure S14:** Co-folding analysis of reported antisense oligonucleotides **(A)** Stem 2 disruptor (S2D), **(B)** Stem 3 disruptor-1 (S3D1), **(C)** Stem 3 disruptor-2 (S3D2), **(D)** Stem 1 disruptor-1, and **(E)** Stem 1 disruptor-3 (S1D3) with 88nt FSE of SARS-CoV-2. Majorly, the binding of antisense oligonucleotides disrupts the SARS-CoV-2 FSE architecture, except for S3D2, where the conformation remains the same.

**Table S1:** Sequence and dot-bracket notation of each viral structure of each construct. Dot bracket notation was generated through IPknot. The red region in the sequences denotes the slippery site for its respective frameshift region

| Virus | 85 | 100-3’ | 5’-100 | 115 |
|---|---|---|---|---|
| **JEV** | CCCUUUUCAGCUGGGC<br>CUUCUGGUGAUGUUU<br>CUGGCCACCCAGGAGG<br>UCCUUCGCAAGAGG<br>UGGACGGCCAGAUUGA<br>CCAUUCCU<br><br>............(((((<br>((((((((.....[[[<br>[[[)).)))))))))))<br>(((....))).....<br>..]]]]]].........<br>... | CCCUUUUCAGCUGGGC<br>CUUCUGGUGAUGUUU<br>CUGGCCACCCAGGAGG<br>UCCUUCGCAAGAGGU<br>GGACGGCCAGAUUGAC<br>CAUUCCUGCGGUUUUG<br>GGGGCC<br><br>............(([[(<br>((((((.......[[[<br>[[[[..)))))))[[))<br>(((....)))....<br>]]]]]]]]]........<br>................<br>]]. | UGGUGAAAUGGUUGAC<br>CCUUUUCAGCUGGGCC<br>UUCUGGUGAUGUUUCU<br>GGCCACCCAGGAGGUCC<br>UUCGCAAGAGGUGGACG<br>GCCAGAUUGACCAUUCCU<br><br>....[[((((((((....<br>...]]....((((((<br>(((((((.....[[[[[[<br>)).)))))))))))<br>(((....))).......]<br>]]]]].))))))))... | UGGUGAAAUGGUUGAC<br>CCUUUUCAGCUGGGCC<br>UUCUGGUGAUGUUUCU<br>GGCCACCCAGGAGGUCC<br>UUCGCAAGAGGUGGACG<br>GCCAGAUUGACCAUUCC<br>UGCGGUUUUGGGGGCC<br><br>....(([[[[[[[[[......)<br>).....(((((<br>((((........[[[[[[[...<br>)))))))))(<br>(((....)))).....]]]]]]<br>]]]]]]]]]]<br>.................. |
| **WNV** | GAUUGAUCCUUUUCAG<br>UUGGGCCUUAUGGUCG<br>UGUUCUUGGCCACCCAG<br>GAGGUCCUUCGCAAGAG<br>GUGGACGGCCAAGAUCA<br>GC<br><br>..((((......)))).<br>..((((....))))..<br>..((((((((.......<br>.(((((((....))))<br>..))))))))))).....<br>.. | GAUUGAUCCUUUUCAG<br>UUGGGCCUUAUGGUCG<br>UGUUCUUGGCCACCCAG<br>GAGGUCCUUCGCAAGAG<br>GUGGACGGCCAAGAUCA<br>GCAUUCCAGCUAUCAUG<br><br>....((......))(((<br>((((((....)))).(<br>(.((((((((.......<br>..(((((((....)))<br>)..))))))))))).))<br>......)))))...... | AUACAACGCCGACAUG<br>AUUGAUCCUUUUCAGU<br>UGGGCCUUAUGGUCGU<br>GUUCUUGGCCACCCAGG<br>AGGUCCUUCGCAAGAGG<br>UGGACGGCCAAGAUCAGC<br><br>..................<br>.((......))....<br>.((((....))))....(<br>(((((((......<br>...(((((((....))))<br>..))))))))))<br>)..... | AUACAACGCCGACAUG<br>AUUGAUCCUUUUCAGU<br>UGGGCCUUAUGGUCGU<br>GUUCUUGGCCACCCAGG<br>AGGUCCUUCGCAAGAGG<br>UGGACGGCCAAGAUCAGC<br><br>............(((((..((.<br>.....))..((<br>(((((....))))....(((((<br>(((.......<br>..((((((((....))))..)))<br>))))))))..<br>[[.....)))]].))))) |
| **HCV** | CCUAAACCUCAAAGAA<br>AAACCAAACGUAACAC<br>CAACCGUCGCCCACAG<br>GACGUCAAGUUCCCGG<br>GUGGCGGUCAGAUCGU<br>UGGUG<br><br>.................<br>...[[.[[[....((<br>..((((((((((...((<br>((.....)).)).<br>))))))))))).....]]<br>]]])). | CCUAAACCUCAAAGAA<br>AAACCAAACGUAACACC<br>AACCGUCGCCCACAGGA<br>CGUCAAGUUCCCGGGUG<br>GCGGUCAGAUCGUUGGU<br>GGAGUUUACUUGUUGC<br><br>.................<br>............(((((<br>((((((((((...((((<br>.....)).)).)))<br>)))))........))))<br>))).............<br>.. | ACCAUGAGCACGAAU<br>CCUAAACCUCAAAGAA<br>AAACCAAACGUAACACC<br>AACCGUCGCCCACAGGA<br>CGUCAAGUUCCCGGGUG<br>GCGGUCAGAUCGUUGGU<br>G<br><br>.....(((........[[<br>....)))..]]...<br>.................(<br>(((((((((...((<br>((.....)).)).)))))<br>)))))........<br>.... | ACCAUGAGCACGAAU<br>CCUAAACCUCAAAGAA<br>AAACCAAACGUAACACC<br>AACCGUCGCCCACAGGA<br>CGUCAAGUUCCCGGGUG<br>GCGGUCAGAUCGUUGGU<br>GGAGUUUACUUGUUGC<br><br>......................<br>..........<br>............((((((((((<br>(((((...<br>((((.....)).)).)))))))<br>)........<br>)))))))............... |

| | | | | |
|---|---|---|---|---|
| **HIV** | GCUAAUUUUUUAGGGA<br>AGAUCUGGCCUUCCUAC<br>AAGGGAAGGCCAGGGAA<br>UUUUCUUCAGAGCAGAC<br>CAGAGCCAACAGCCCCA<br>C<br><br>.................<br>...[[.[[[....((.<br>.((((((((((...(((<br>(.....)).)).))<br>))))))))).....]]]]<br>])). | GCUAAUUUUUUAGGG<br>AAGAUCUGGCCUUCCU<br>ACAAGGGAAGGCCAGG<br>GAAUUUUCUUCAGAGC<br>AGACCAGAGCCAACAGC<br>CCCACCAGAAGAGAGCU<br>UCA<br><br>.((((....))))....<br>...(((((((((((<br>....))))))))))).. <br>...[[[.........<br>.................<br>.......((((]]].))<br>)). | UGUACUGAGAGACAGG<br>CUAAUUUUUUAGGGAA<br>GAUCUGGCCUUCCUACA<br>AGGGAAGGCCAGGGAAU<br>UUUCUUCAGAGCAGACCA<br>GAGCCAACAGCCCCAC<br><br>.[[...........(((.<br>.........(((((<br>..((((((((((((....<br>))))))))))))<br>[[[)))))]]].......<br>..........]].)<br>)).... | UGUACUGAGAGACAGG<br>CUAAUUUUUUAGGGAA<br>GAUCUGGCCUUCCUACA<br>AGGGAAGGCCAGGGAAU<br>UUUCUUCAGAGCAGACCA<br>GAGCCAACAGCCCCACCAGAAG<br>AGAGCUUCA<br><br>................[[....<br>...........<br>.((((((((((((....)))))<br>)))))))<br>......................<br>............<br>......((((..]])))). |

**Table S2:** Non-canonical interactions stabilizing the 85nt, 100-3'nt, 5'-100nt, and 115nt constructs for JEV, WNV, HCV, and HIV.

| | | |
|---|---|---|
| **85nt** | | |
| JEV | simRNA | G15-U48, G60-U51, U19-G44, G14-U62-G14, G61-G13-C50, U12-G58-C53 |
| HIV | | A61-A12-U3, U57-G13, U56-G14, G18-U52, A71-C81, A69-C83 |
| HCV | | A4-C10, U27—A28-U77, C33-A74, A34-C73, U39-G68, G41-U66, A45-C62, A47-A46-G49 |
| WNV | | G60-A63, G16-U3, G1-U18, U30-G21, G84-U33, G19-G32-C85 |
| JEV | trRosettaRNA | C32-U33-G72, U27-C70-G25, G67-C40-G23, U19-G44, G15-U48, A9-G54, U5-G58, G60-U4-A59 |
| HIV | | U31-G36, G47-U20, G48-A19, U52-G14, G13-U53 |
| HCV | | A47-C61, G41-U66, G68-U39 |
| WNV | | G73-A44, C42-U30, G32-A71 |
| **5'-100nt** | | |
| JEV | simRNA | G59-U34, G30-U63, G75-U66, G91-U13 |
| HIV | | G51-U46, G62-U35, G63-A34, U68-G30, U69-G29, G75-U21, G7-A91-U3 |
| HCV | | A71-G67-A70, G80-A60, U81-G56, G83-U54, A71-G67-A70 |
| WNV | | U82-G71, U22-G67, U33-G16, U18-G31 |
| JEV | trRosettaRNA | U63-G30, G69-A24, G82-U13, G83-U12, G87-A8, G91-G5 |
| HIV | | G51-U46, G62-U35, G65-A33, A4-A76 |
| HCV | | A28-C3, A39-G94-C40, G41-U92, A49-C88, C47-A89, U54-G83, U81-G56, A60-C77, G67-A71 |
| WNV | | U19-U27-U18, U33-G11, G36-G8, U50-U96, A78-G25 |
| **100-3'nt** | | |
| JEV | simRNA | G60-U51, G15-U48, U19-G44, G76-C100-G28, U81-G96, G92-G86 |
| HIV | | G65-A64, G62-A66, A59-A69, G93-U53, G95-U51, G91-C96-A50, C86-A19, A78-A16, G1-G14-C82, G36-U31, A75-G47, G91-A49-A92 |
| HCV | | C62-A45, U66-G41, G68-U39, C73-A34, C33-A74, A4-U9-A5 |
| WNV | | G92-U17, U30-G21 |
| JEV | trRosettaRNA | U93-U4, U92-U5, U91-U6, U90-U7, C16-A80, C17-C79, G54-A57 |
| HIV | | G36-U31, G47-U20, G15-A84, G93—G91-U7 |
| HCV | | G41-U66, G68-U39 |
| WNV | | A63-G60, U68-U57, A71-C55, U81-U35 |
| **115nt** | | |
| JEV | | U45-G49, U44-G50, U34-G59, U63-G30, G75-U66, G69-A72, G113-U12, G14-G111-C16, U100-U77, C47-G49-U45 |

| | | |
|---|---|---|
| HIV | simRNA | C96-A86, C83-A19, G80-U22, A73-U29, A79-U23, U24-G77, U25-G75, G28-U72, G29-U71, G33-U67, A34-A65, U35-G62, G51-U46 |
| HCV | | A10-A20, A27-A107-A28, A1-U109-A30, A31-U106-U112, U105-A32-U106, U104-C35, G94-C52-G86, G83-U54, G56-U81, C61-C77, G67-A71 |
| WNV | | A6-U22-U27, C7-A21-U88, C45-G36, G47-U103, U48-G99, G75-A78 |
| JEV | trRosettaRNA | U100-G2, U13-G91, G19-U90, U89-U45, G30-U63, U66-G76, A86-G49 |
| HIV | | U46-G51, U35-G62, U69-U25, A12-C13-G2 |
| HCV | | U74-A65, G56-U81, G83-U54 |
| WNV | | U83-U72, A78-G75, U50-U96 |

**Table S3:** Information about ASO sequences generated from Oligowalk and their scrambled counterpart.

| JEV 85 | | | | | | | | | | |
|---|---|---|---|---|---|---|---|---|---|---|
| 21nt (ASO) | | | | | | | | | | Scrambled ASO |
| Pos. | Oligo(5'->3') | Overall | Duplex | Tm-Dup | Break-targ. | Intraoligo | Interoligo | End_diff | prefilter_score | |
| kcal/mol | kcal/mol | kcal/mol | kcal/mol | kcal/mol | kcal/mol | | | | | |
| 10 | AACAUCACCAGAAGGCCCAGC | -26.1 | -42.2 | 92.7 | -12.8 | -0.9 | -14.7 | 2.49 | 6 | **CCACGUGCAGACGACACAAAC** |
| 54 | AUCUGGCCGUCCACCUCUUGC | -29.1 | -44 | 94.3 | -10.7 | -1.3 | -16.5 | 2.32 | 6 | **UCGGUUUCCCCCAUGGAUCCC** |
| **JEV 115** | | | | | | | | | | |
| **21nt (ASO)** | | | | | | | | | | **Scrambled ASO** |
| Pos. | Oligo(5'->3') | Overall | Duplex | Tm-Dup | Break-targ. | Intraoligo | Interoligo | End_diff | prefilter_score | |
| kcal/mol | kcal/mol | kcal/mol | kcal/mol | kcal/mol | kcal/mol | | | | | |
| 25 | AACAUCACCAGAAGGCCCAGC | -25.5 | -42.2 | 92.7 | -13.4 | -0.9 | -14.7 | 2.49 | 6 | **CCCACGAAUGCACGCCAAAGA** |
| 86 | AACCGCAGGAAUGGUCAAUCU | -24.6 | -37.6 | 87.5 | -9.9 | -2.7 | -13.9 | 0.7 | 6 | |
| **WNV 85** | | | | | | | | | | |
| **21nt (ASO)** | | | | | | | | | | **Scrambled ASO** |
| Pos. | Oligo(5'->3') | Overall | Duplex | Tm-Dup | Break-targ. | Intraoligo | Interoligo | End_diff | prefilter_score | |
| kcal/mol | kcal/mol | kcal/mol | kcal/mol | kcal/mol | kcal/mol | | | | | |
| 8 | CAUAAGGCCCAACUGAAAAGG | -23 | -36.6 | 86.6 | -10.3 | -0.8 | -14.7 | 0.7 | 6 | **GACAACAGUAGCCUAACAGAG** |
| 19 | AAGAACACGACCAUAAGGCCC | -21.9 | -39.3 | 89.9 | -14.2 | -0.8 | -14.6 | 2.33 | 6 | **GCGCACCAAAACAUAACGCGA** |
| **WNV 115** | | | | | | | | | | |
| **21nt (ASO)** | | | | | | | | | | **Scrambled ASO** |
| Pos. | Oligo(5'->3') | Overall | Duplex | Tm-Dup | Break-targ. | Intraoligo | Interoligo | End_diff | prefilter_score | |

| | | | | | | | | | | |
|---|---|---|---|---|---|---|---|---|---|---|
| kcal/mol | kcal/mol | kcal/mol | kcal/mol | kcal/mol | kcal/mol | | | | | |
| 2 | AUCAAUCAUGUCGGCGUUGUA | -24.9 | -35 | 83.7 | -5.1 | -0.5 | -18.1 | 0.23 | 7 | **GUCAGUGGCUCAAUUUAUGAC** |
| 34 | AAGAACACGACCAUAAGGCCC | -24.7 | -39.3 | 89.9 | -11.4 | -0.8 | -14.6 | 2.33 | 6 | |
| **HCV 85** | | | | | | | | | | |
| **21nt (ASO)** | | | | | | | | | | **Scrambled ASO** |
| Pos. | Oligo(5'->3') | Overall | Duplex | Tm-Dup | Break-targ. | Intraoligo | Interoligo | End_diff | prefilter_score | |
| kcal/mol | kcal/mol | kcal/mol | kcal/mol | kcal/mol | kcal/mol | | | | | |
| 7 | ACGUUUGGUUUUUCUUUGAGG | -27.9 | -31.4 | 80.4 | -1.5 | -0.5 | -11.9 | 1.47 | 6 | **GUUGAUGCGUUUCUGUAUUUG** |
| 39 | AACUUGACGUCCUGUGGGCGA | -18.8 | -41.3 | 92.6 | -16 | -6 | -20.8 | 0.97 | 6 | **ACGGUAGGCUACUGUUCUAUA** |
| **HCV 115** | | | | | | | | | | |
| **21nt (ASO)** | | | | | | | | | | **Scrambled ASO** |
| Pos. | Oligo(5'->3') | Overall | Duplex | Tm-Dup | Break-targ. | Intraoligo | Interoligo | End_diff | prefilter_score | |
| kcal/mol | kcal/mol | kcal/mol | kcal/mol | kcal/mol | kcal/mol | | | | | |
| 1 | UUUAGGAUUCGUGCUCAUGGU | -30.6 | -35.6 | 85.1 | -2.3 | -0.7 | -13.5 | 0.86 | 7 | **AGUGCAUUGUGGUAUUGCUUC** |
| 86 | AAACUCCACCAACGAUCUGAC | -23.1 | -37.1 | 86 | -12.1 | -0.2 | -11.9 | 1.76 | 7 | |
| **HIV 85** | | | | | | | | | | |
| **21nt (ASO)** | | | | | | | | | | **Scrambled ASO** |
| Pos. | Oligo(5'->3') | Overall | Duplex | Tm-Dup | Break-targ. | Intraoligo | Interoligo | End_diff | prefilter_score | |
| kcal/mol | kcal/mol | kcal/mol | kcal/mol | kcal/mol | kcal/mol | | | | | |
| 1 | GAUCUUCCCUAAAAAAUUAGC | -22 | -30.5 | 77.7 | -6.7 | -0.7 | -11.7 | 0.62 | 6 | **AUGACUAAUCAUCACUUACGA** |
| 32 | AAUUCCCUGGCCUUCCCUUGU | -16.8 | -40.3 | 92 | -20 | -0.6 | -15.1 | 0.86 | 6 | **UUUACCCCUCUAGUCGCUCGU** |

| HIV 115 | | | | | | | | | | |
|---|---|---|---|---|---|---|---|---|---|---|
| 21nt (ASO) | | | | | | | | | | Scrambled ASO |
| Pos. | Oligo(5'->3') | Overall | Duplex | Tm-Dup | Break-targ. | Intraoligo | Interoligo | End_diff | prefilter_score | |
| kcal/mol | kcal/mol | kcal/mol | kcal/mol | kcal/mol | kcal/mol | | | | | |
| 1 | AUUAGCCUGUCUCUCAGUACA | -29.7 | -36.8 | 86.9 | -4.9 | -1.8 | -12.3 | 0.56 | 8 | **UCAACGUUCGAACAUUGUCUC** |
| 47 | AAUUCCCUGGCCUUCCCUUGU | -18 | -40.3 | 92 | -18.8 | -0.6 | -15.1 | 0.86 | 6 | |

**Table S4:** Negative △G between FSE-ASO heteroduplex generated from RNA co-fold analysis

| Viral Frameshifting Element | Length | △G of FSE-ASO heteroduplex (kcal/mol) | |
|---|---|---|---|
| | | **ASO 1** | **ASO 2** |
| **JEV** | 85 nt | -30.95 | -34.03 |
| **WNV** | 85 nt | -28.18 | -27.16 |
| **HCV** | 85 nt | -31.03 | -13.14 |
| **HIV** | 85 nt | -24.50 | -19.74 |
| **JEV** | 115 nt | -28.20 | -24.77 |
| **WNV** | 115 nt | -31.50 | -27.04 |
| **HCV** | 115 nt | -34.32 | -26.46 |
| **HIV** | 115 nt | -31.48 | -20.72 |
